# Extensive transmission and variation in a functional receptor for praziquantel resistance in endemic *Schistosoma mansoni*

**DOI:** 10.1101/2024.08.29.610291

**Authors:** Duncan J. Berger, Sang-Kyu Park, Thomas Crellen, Tushabe John Vianney, Narcis B. Kabatereine, James A. Cotton, Richard Sanya, Alison Elliot, Edridah M. Tukahebwa, Moses Adriko, Claire J. Standley, Anouk Gouvras, Safari Kinung’hi, Helmut Haas, Muriel Rabone, Aidan Emery, Poppy H. L. Lamberton, Bonnie L. Webster, Fiona Allan, Sarah Buddenborg, Matthew Berriman, Jonathan S. Marchant, Stephen R. Doyle, Joanne P. Webster

**Author notes:** Corresponding authors Underline. co-supervised this work (Matthew Berriman, Jonathan S. Marchant, Stephen R. Doyle, Joanne P. Webster).

## Abstract

Mass-drug administration (MDA) of human populations using praziquantel monotherapy has become the primary strategy for controlling and potentially eliminating the major neglected tropical disease schistosomiasis. To understand how long-term MDA impacts schistosome populations, we analysed whole-genome sequence data of 570 *Schistosoma mansoni* samples (and the closely related outgroup species, *S. rodhaini)* from eight countries incorporating both publicly-available sequence data and new parasite material. This revealed broad-scale genetic structure across countries but with extensive transmission over hundreds of kilometres. We characterised variation across the transient receptor potential melastatin ion channel, TRPM_PZQ_, a target of praziquantel, which has recently been found to influence praziquantel susceptibility. Functional profiling of TRPM_PZQ_ variants found in endemic populations identified four mutations that reduced channel sensitivity to praziquantel, indicating standing variation for resistance. Analysis of parasite infrapopulations sampled from individuals pre- and post-treatment identified instances of treatment failure, further indicative of potential praziquantel resistance. As schistosomiasis is targeted for elimination as a public health problem by 2030 in all currently endemic countries, and even interruption of transmission in selected African regions, we provide an in-depth genomic characterisation of endemic populations and an approach to identify emerging praziquantel resistance alleles.

**One Sentence Summary:** Population genomics and functional genetics of praziquantel resistance in *Schistosoma mansoni*

## INTRODUCTION

Schistosomiasis is a neglected tropical disease which currently infects 250 million people across 78 endemic nations (*1*, *2*). Infections are prevalent among children, including pre-school-aged children, as well as adults in low and middle-income countries, with 90% of infections occurring in sub-Saharan Africa. The etiological agents of schistosomiasis are freshwater snail-borne parasitic trematodes of the genus *Schistosoma* (principally *Schistosoma mansoni, S. japonicum,* and *S. haematobium*). These parasitic worms dwell in the host’s blood vessels, often for years, where they lay eggs, many of which become trapped in host tissues, resulting in a spectrum of pathologies including anaemia, stunted growth, genital lesions, fever and irreversible organ damage (*3*). Recognising the severe and widespread impact of this disease, the World Health Organization (WHO) endorsed praziquantel monotherapy, distributed as part of mass drug administration (MDA) programmes, as the primary strategy for schistosomiasis control (*4*, *5*). In 2020 alone, 76.9 million people (representing 44.9% of those requiring treatment), predominantly school-aged children, were treated with praziquantel for schistosomiasis (*6*). Such treatment campaigns have resulted in an overall decrease in the prevalence of schistosomiasis among school-aged children by approximately 60% (*7*, *8*). Based on such successes, the WHO launched its 2021-2030 NTD roadmap and revised Guidelines for the Control and Elimination of Schistosomiasis (*1*). These set ambitious goals to eliminate schistosomiasis as a public health problem in all endemic countries (defined as reducing the proportion of heavy-intensity infections to <1%) and complete interruption of transmission in selected regions by 2030, primarily through the escalating use of praziquantel MDA (*9*).

Although MDA programmes have, in general, resulted in large-scale reductions in schistosomiasis prevalence and morbidity in endemic regions (*7*, *10*, *11*), some of these prevalence reductions have proven to be reversible over short timescales and, despite years of repeated MDA, persistent hotspots of infection remain (*12–15*). Ongoing surveillance projects have also revealed substantial heterogeneity in infection prevalence, intensity and morbidity within and between endemic regions (*16–18*). This includes potential variability in parasite response to praziquantel, which, given the current lack of available vaccines or alternative antischistosomal drugs, represents a major threat to the control and elimination of schistosomiasis (*19–21*). In combination with other public health measures, MDA programmes are expected to dramatically alter patterns of schistosome transmission and exert selective pressure for praziquantel resistance. However, despite over twenty years of MDA in some areas, there is no consistent evidence that sustained praziquantel administration has impacted parasite population genetic structure or diversity (*22*–*25*). Likewise, while there is clear evidence that resistance to praziquantel can be rapidly selected for in laboratory settings (*26–29*), there is limited evidence of established praziquantel resistance in endemic *S. mansoni* populations (*30*). This contrasts with that of emerging or established heritable resistance, particularly in the veterinary helminths, to every other class of anthelmintic used (*31–34*). Part of the explanation lies in the fact that, until recently, we neither knew the precise mode of action of praziquantel against schistosomes nor had any molecular markers to monitor potential praziquantel resistance amongst natural *Schistosoma* spp. populations. Recent efforts have, for the first time, identified the molecular target of praziquantel, a transient receptor potential (TRP) melastatin ion channel (*Sm*.TRPM_PZQ_) (*35–38*). This discovery permits evaluation of praziquantel efficacy and resistance risk, although such resistance-conferring alleles have only been identified in a single isolate from endemic populations (*37*). Characterising the transmission and recent evolution of schistosome populations is, therefore, of broad epidemiological importance as a means to understand how parasite populations are structured and how they are changing in response to interventions aimed at controlling schistosomiasis.

Here, we have used whole-genome sequencing to characterise genomic variation from globally dispersed populations of the major human infective species *S. mansoni* (and the closely related outgroup species, *S. rodhaini)* (*39–41*). Our analyses focus on populations within endemic regions of East Africa, primarily those found around Lake Victoria, one of the largest foci of schistosomiasis infections and a target of long-term MDA efforts (*7*, *20*). We aimed to quantify variation within the newly identified candidate praziquantel resistance locus, *Sm*.TRPM_PZQ_ (*36*, *37*, *42*), to determine the prevalence of potential resistance-conferring mutations. This also included extensive WGS of *S. mansoni* from within individual hosts before and after praziquantel administration to quantify the immediate genetic impact of treatment. These data raise significant implications regarding the standing variation of potential praziquantel resistance in human schistosome population and represent an important resource for assessing the efficacy of current interventions and guiding future treatment strategies.

## RESULTS

### The genetic diversity of geographically dispersed isolates supports high transmission in endemic regions

We analysed whole-genome sequence data from 574 *Schistosoma* samples (*n* = 570 *S. mansoni* and *n* = 4 *S. rodhaini*; hereafter all referred to as accessions) isolated from eight countries, including 207 new *Schistosoma* samples sequenced for this study (Fig. 1A; Table 1; Supplementary Data 1) (*39–41*). Most accessions were derived from Lake Victoria (88.3%), a major focus of *S. mansoni* infection.

**Fig. 1:**
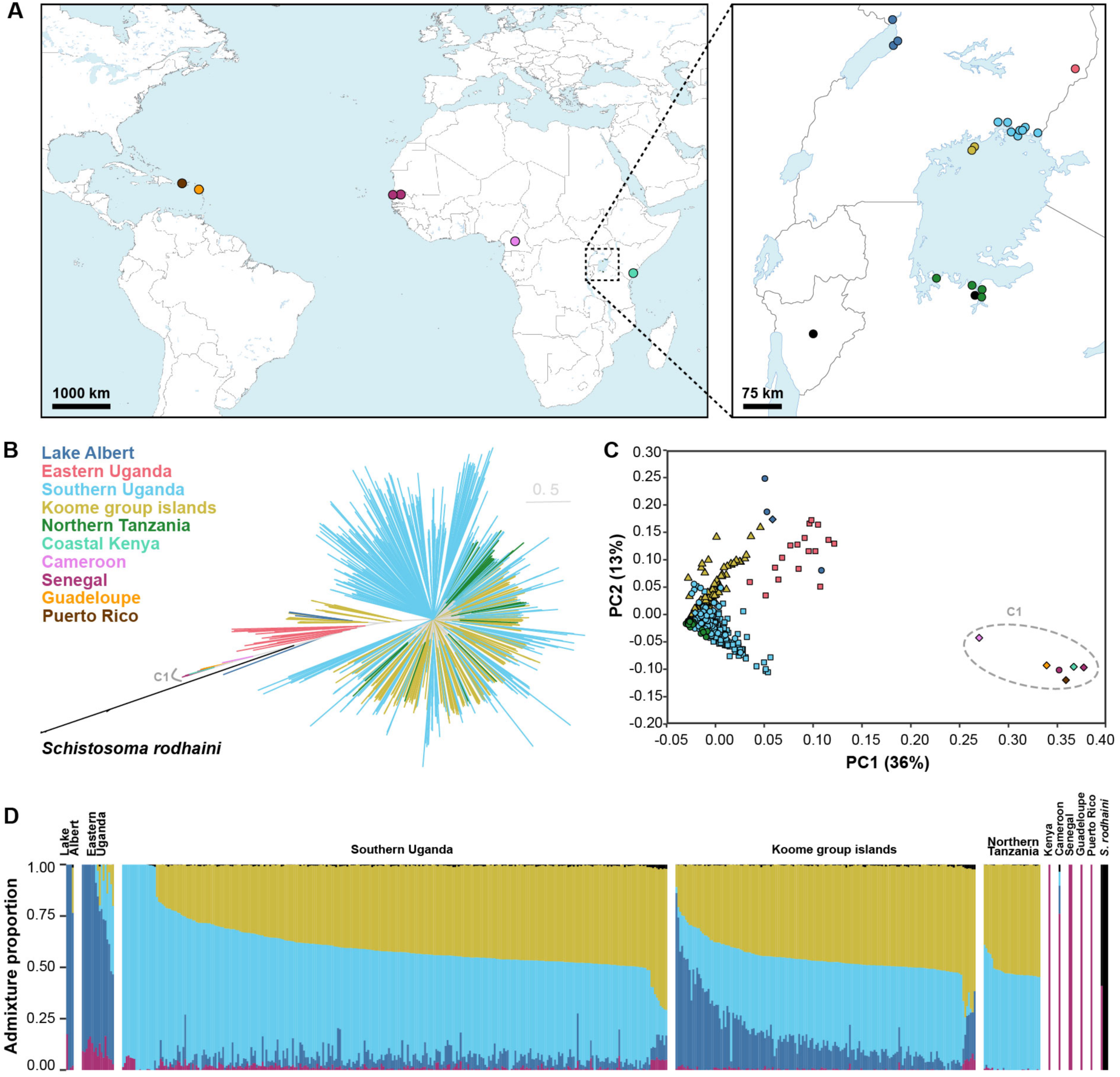
Population structure of *Schistosoma* accessions. **A.** Global distribution of the 574 *Schistosoma* accessions used in this study. Sampling locations are coloured by geographical divisions: Puerto Rico (PR; brown), Guadeloupe (GP; orange), Senegal (SN; dark purple), Cameroon (pink), Coastal Kenya (KE; teal), Lake Albert (dark blue), Eastern Uganda (red), Southern Uganda (light blue), Koome group islands (yellow), Northern Tanzania (dark green). Black points represent the sampling coordinates of *S. rodhaini* accessions. **B.** The maximum-likelihood phylogenetic tree was inferred using 188,923 autosomal single-nucleotide polymorphisms (SNPs) and all 574 accessions. Branches are coloured based on the subdivisions described in a), and the tree is rooted on *S. rodhaini*. Highlighted clade ‘C1’ represents all West African (Senegalese and Cameroonian), Caribbean (Puerto Rican and Guadeloupean) and Coastal Kenyan accessions. **C.** Principal component analysis (PCA) of genetic differentiation between 505 unrelated *S. mansoni* accessions using 214,445 autosomal SNPs. Points are coloured and shaped based on the groups described in (A). **D.** ADMIXTURE plots illustrating the inferred ancestry of 505 unrelated *S. mansoni* and four *S. rodhaini* accessions. Here, we assume five populations (K) are present, inferred using 10-fold cross-validation and a standard error estimation with 250 bootstraps. Y-axis values show the admixture proportions for each accession, and colours for each population were assigned based on the majority ancestry of each geographical division.

**Table 1:** Sample information and history of praziquantel treatment.

| Species | Country | Region | Development Stage |  |  | Sampling stage |  |  | Reference |
| --- | --- | --- | --- | --- | --- | --- | --- | --- | --- |
|  |  |  | Adults | Cercariae | Miracidia | Pre-treatment | Post-treatment | Not applicable |  |
| *S. mansoni* | Uganda | Lake Albert | 1 | 6 | 0 | 0 | 0 | 7 | Herein; Crellen et al. 2016 |
|  | Uganda | Eastern Uganda | 0 | 0 | 17 | 17 | 0 | 0 | Berger et al. 2021 |
|  | Uganda | Southern Uganda | 1 | 2 | 328 | 201 | 127 | 3 | Herein; Berger et al. 2021 |
|  | Uganda | Koome Islands | 0 | 0 | 174 | 123 | 51 | 0 | Vianney et al. 2022 |
|  | Tanzania | Northern Tanzania | 0 | 31 | 0 | 0 | 0 | 31 | Herein |
|  | Kenya | Coastal Kenya | 1 | 0 | 0 | 0 | 0 | 1 | Crellen et al. 2016 |
|  | Cameroon |  | 1 | 0 | 0 | 0 | 0 | 1 | Crellen et al. 2016 |
|  | Senegal |  | 3 | 0 | 0 | 0 | 0 | 3 | Herein; Crellen et al. 2016 |
|  | Puerto Rico |  | 1 | 0 | 0 | 0 | 0 | 1 | Crellen et al. 2016 |
|  | Guadeloupe |  | 4 | 0 | 0 | 0 | 0 | 4 | Crellen et al. 2016 |
| *S. rodhaini* | Tanzania | Northern Tanzania | 0 | 2 | 0 | 0 | 0 | 2 | Herein |
|  | Burundi |  | 2 | 0 | 0 | 0 | 0 | 2 | Herein; Crellen et al. 2016 |

This study included 574 accessions, sequenced from parasite samples derived from snail hosts (cercariae), humans (miracidia) or laboratory-
passaged strains (adults). Publicly-available data ( $n = 339$ ) and new sequencing data ( $n = 208$ ; defined as “herein”) were generated for this
study. Miracidia accessions were sampled pre-treatment with praziquantel (40 mg/kg) or four weeks post-treatment.

These included published sequence data from miracidial samples isolated from infected children between 2014 and 2017 from either the Koome & Damba Islands (hereafter referred to as the Koome Islands; *n =* 174) (*41*) or Southern Uganda (*n =* 164) (*39*), with a median of two isolates sequenced per child (range 1-11). We sequenced an additional 164 miracidia from just three children from Southern Uganda (*n =* 89, 51 & 23 miracidia per child), including both pre-(*n* = 82) and post-praziquantel treatment (*n* = 82) sampling (*20*). The only other Lake Victoria accessions were derived from cercariae shed from snails captured in Northern Tanzania (*n =* 31) or Southern Uganda (*n* = 2). Of the remaining samples, two were cercariae from the shoreline of Lake Albert, four were cercariae from a passaged strain originally isolated in Lake Albert, 17 were miracidia collected from children living in Eastern Uganda, and ten were published adult-stage isolates of passaged laboratory strains originally sampled from Kenya (*n =* 1), Senegal (*n =* 3), Cameroon (*n* = 1), Guadeloupe (*n =* 4) or Puerto Rico (*n =* 1) (*40*). As an outgroup, we included four *S. rodhaini* accessions, two cercarial isolates from Tanzania and two adult-stage isolates of passaged lab strains from isolates originally sampled in Burundi. We mapped sequence reads from all 574 accessions to the *S. mansoni* reference genome, and after variant calling and quality control, we identified 35,146,249 single-nucleotide polymorphisms (SNPs) and 6,632,156 indels across all accessions (Supplementary Data 2).

To characterise population structure, we produced a maximum likelihood phylogeny and performed principal component analysis using unlinked autosomal variants. These analyses distinguished accessions sampled in East Africa from those in West Africa and the Caribbean (Fig. 1B,C). The only exception was a single laboratory-passaged isolate from coastal Kenya that clustered with West African accessions, consistent with previous analyses of this sample (*40*, *43*). Within Eastern Africa, populations from Lake Victoria, Lake Albert and Eastern Uganda formed distinct but partially overlapping clusters. Pairwise measures of the fixation index (*F*_ST_) showed limited population differentiation between Lake Victoria, Lake Albert and Eastern Uganda (range *F*_ST_ = 2.49×10^-2^ - 4.59×10^-2^) and negligible differentiation between Lake Victoria populations, despite the geographic separation of up to 300 km (range *F*_ST_ = 2.41×10^-3^ - 4.07×10^-3^, Supplementary Fig. 1A). East African populations had comparable levels of nucleotide diversity (range median π = 2.35×10^-3^ - 3.35×10^-3^; Supplementary Fig. 1B) and effective population size (*N_e_*) estimates (range *N_e_* = 58,004-64,063; Supplementary Fig. 2; Supplementary Data 3) suggesting shared recent population histories. Analyses of population ancestry for individual accessions identified similar population compositions across all Lake Victoria populations with variable ancestry contributions from Lake Albert and West African populations (Fig. 1F, Supplementary Data 4). Accessions from Eastern Uganda displayed variable, mixed ancestry from Lake Victoria, Lake Albert and West Africa populations, suggesting migration from multiple regions.

Ancestry analyses revealed low levels of *S. rodhaini* admixture (range: 0.00-2.32%), potentially indicative of interspecific hybridisation, across most unrelated *S. mansoni* accessions (Fig. 1D; Supplementary Data 4). In East Africa, *S. mansoni* and *S. rodhaini* are broadly sympatric and share both intermediate and definitive hosts, with *S. mansoni* infecting both specific primate and rodent species and *S. rodhaini* infecting rodents, allowing opportunities for inter-species pairings and hybridisation. However, due to the small number of *S. rodhaini* isolates included in this study (*n* = 4) and the lack of a non-admixed, outgroup population, the incidence of recent admixture and/or historic introgression was not investigated further.

### Functionally relevant genomic variation influences praziquantel sensitivity

A transient receptor potential (TRP) melastatin ion channel (*Sm.*TRPM_PZQ_; *Smp_246790*) has recently been proposed as the target of praziquantel (*35–38*). This candidate gene underlies variation in praziquantel susceptibility, at least within schistosome laboratory populations (*35–37*, *44*). Mutagenesis of key residues within the praziquantel binding site of TRPM_PZQ_ (located in the transmembrane voltage-sensor-like domain, VSLD) resulted in the loss of sensitivity to praziquantel (*36*).

Consistent with previous genomic surveys (*39*, *41*), analysis of haplotype diversity using the integrated haplotype score (iHS) did not suggest that *Sm.*TRPM_PZQ_ was under positive selection in Lake Victoria populations (Supplementary Fig. 3, Supplementary Data 5,6). However, analysis of variation within *Sm.*TRPM_PZQ_ identified a high degree of variability with a predicted 496 amino acid changes at 433/2268 residues (Fig. 2), with mutations at 174 of these residues found only in single accessions (Supplementary Data 7). We generated point mutations to modify 12 *Sm*.TRPM_PZQ_ residues, and used an *in vitro* Ca^2+^ reporter assay (*35*) to assess their impact on *Sm*.TRPM_PZQ_ function. Eight of these mutants, all conservative amino acid substitutions, exhibited similar praziquantel sensitivity (range EC_50_ = 0.49-0.96 *µ*M) to the wild-type *Sm*.TRPM_PZQ_ channel (EC_50_ = 0.74 *µ*M [standard error (SE) = 0.17]; Fig. 3A; Supplementary Data 8; Supplementary Fig. 4). Mutants p.T1624K and p.R1843Q exhibited a decreased sensitivity to praziquantel (EC_50_ = 1.41 *µ*M [SE = 0.23] and EC_50_ = 1.11 *µ*M [SE = 0.08], respectively) and two mutants (p.Y1554C, p.Q1670K) caused a complete loss in praziquantel sensitivity (Fig. 3B; Supplementary Data 8; Supplementary Fig. 5).

**Fig. 2:**
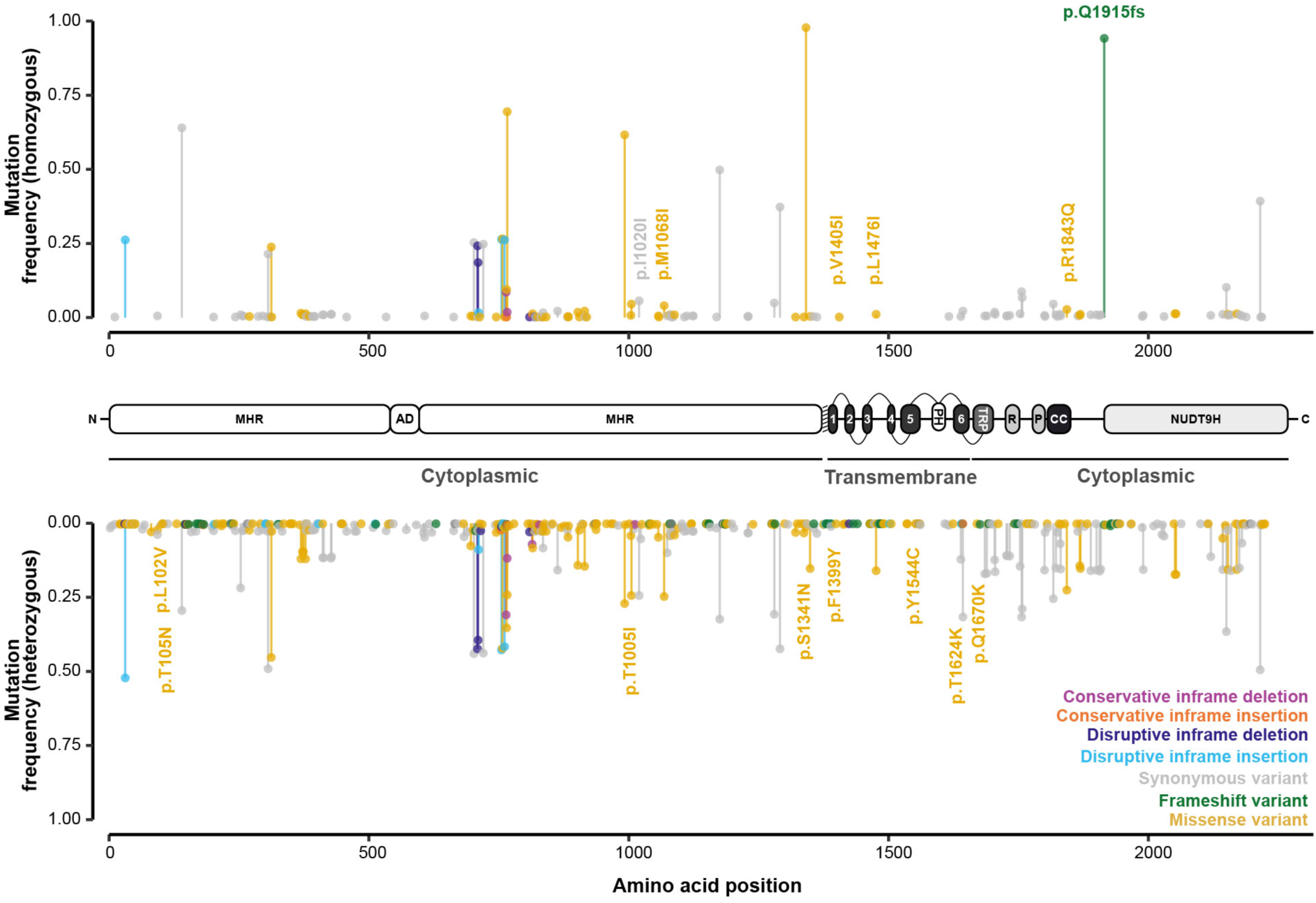
Genetic variation in the candidate mediator of praziquantel susceptibility, the transient potential receptor channel *Sm.*TRPM_PZQ_. Mutational frequency along the protein structure of *Sm.*TRPM_PZQ_ (*Smp_246790*). Frequencies of mutations are reported across 550 *S. mansoni* accessions (y-axis), representing samples from Eastern Uganda (*n* = 17), Southern Uganda (*n* = 328), the Koome islands (*n* = 174) and Northern Tanzania (*n* = 31). X-axis values represent the location of the mutations on the protein, and bars (and terminal points) are coloured by the predicted impact of each mutation. Structure of *Sm.*TRPM_PZQ_: N-terminal TRPM homology region (MHR) domain, ankyrin-like repeat domain (AD), pre-S1 helix (shaded), TM-spanning helices (1–6), pore helices (PH), TRP domain (TRP), rib helices (R), pole helices (P), coiled-coil (CC) region and the COOH terminal NUDT9H domain (NUDT9H).

**Fig. 3:**
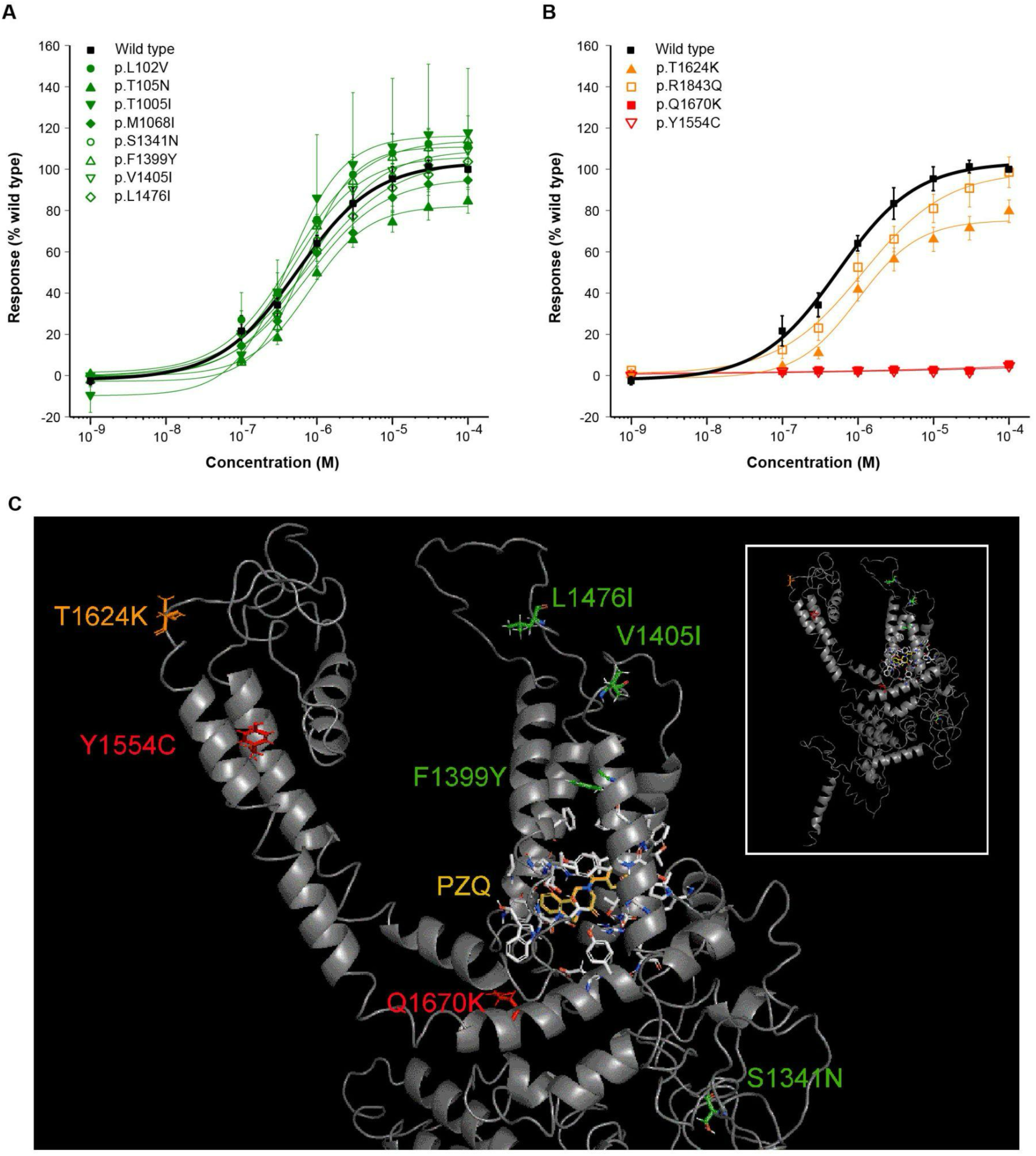
Functional profiling of variants of the transient potential receptor channel *Sm*.TRPM_PZQ_. Concentration-response relationships for consensus Sm.TRPM_PZQ_ sequence compared with twelve Sm.TRPM_PZQ_ variants, which exhibit an average EC_50_ to ±PZQ of **(A)** <1 μM (green) or **(B)** >1 μM (orange). Constructs for which no praziquantel-evoked activity was observed (Y1554C, Q1670K) are shown in (b) in red. Results represent mean±sem from at least three independent transfections. **C.** *Inset*, homology model of the transmembrane spanning region (residues 1100 to 1800) of a *Sm*.TRPM_PZQ_ monomer from [2]. The enlarged view shows the location of the functionally profiled variants. Praziquantel and residues within 5Ǻ of the praziquantel binding poise (white) are shown at the base of the voltage-sensor-like domain (VSLD) of the channel, with the S5 and S6 pore-forming helices to the left.

Seven of the twelve mutations mapped within an existing homology model for the transmembrane-spanning region of *Sm*.TRPM_PZQ_, enabling structural insight into these functional effects (Fig. 3C) (*36*). Four mutations had no impact on praziquantel action (p.V1405I, p.L1476I, p.S1341N & p.F1399Y; Fig. 3A): p.V1405I and p.L1476I are found within extracellular loops of the channel, p.S1341N lies the COOH-terminal cytoplasmic domain and p.F1399Y, while within the voltage-sensor like domain (VSLD) of the channel that harbours the praziquantel binding site, is remote (∼10Ǻ away) from the praziquantel binding poise (Fig. 3C). Mutation p.T1624K (∼2-fold lower sensitivity; Fig. 3B) projects into the extracellular milieu before the start of S6 and may cause sub-optimal orientation of the S6 helix. The other variant associated with lower sensitivity (p.R1843Q; Fig. 3B) lies outside the existing TRPM_PZQ_ homology model but is predicted to localise within the COOH-terminal coiled-coil region of the channel, which is involved in subunit interaction (*45*). Tetramer misfolding and lower channel expression would explain the decreased sensitivity of this variant.

The first of the two mutations that caused a complete loss of praziquantel sensitivity (Fig. 3B), p.Y1554C, is found in the S5 helix (Fig. 3C). This residue projects toward S6, where it is predicted to interact with a tyrosine residue (Y1636) in the S6 helix. This cysteine mutant would be expected to disrupt this interaction, which is likely important for the appropriate orientation of the helices that form the pore-forming domain. The second, p.Q1670K, introduces a positive charge within the intracellular TRP helix close to the bilayer interface. This is a critical region for channel gating, and the mutation likely impacts the orientation of the TRP helix relative to the bilayer and interactivity with the S4/S5 loop of *Sm*.TRPM_PZQ_, which regulates TRPM channel activation (*46*). The deleterious effects of p.Y1554C and p.Q1670K are, therefore, consistent with the known importance of these regions for channel function (*36*).

Of the four mutations that decreased or eliminated praziquantel sensitivity, three (p.Y1554C, p.Q1670K, p.T1624K) were only found in a heterozygous state in single accessions (Supplementary Data 7,9,10). p.R1843Q was far more common, with homozygous variants (c.5528G>A) identified in 3.68% (*n =* 21 accessions across the entire dataset; *n* = 15 miracidia from 12 individuals) of accessions and heterozygous in 22.6% of accessions (*n =* 129 accessions across the entire dataset; *n* = 120 miracidia from 68 individuals). This variant was also more prevalent in post-treatment populations (5.05% of post-treatment accessions; *n* = 9 miracidia from 8 individuals) compared to pre-treatment populations (1.75% of pre-treatment accessions; *n* = 6 miracidia from 4 individuals; Supplementary Data 10), although accessions with this variant (and all other evaluated variants) did not form a distinct subpopulation (Supplementary Fig.s 6-9). In addition to the mutations analysed via targeted mutagenesis, we identified a second high-prevalence homozygous frameshift mutation, p.Q1915fs, in 94.0% of accessions. Park and colleagues (*36*) reported that truncation mutations (Δ1914) of the C-terminal NUDT9H domain do not impact the TRP channels’ responsiveness to praziquantel.

Le Clec’h and colleagues recently identified three putative marker variants for praziquantel resistance in selected laboratory lines (*37*). The first of these was a homozygous SNP predicted to result in a synonymous mutation (*Sm.*TRPM_PZQ_-2723187C; p.I1020l), which we identified in 6.14% (*n =* 35, *n* = 30 miracidia from 21 individuals) of accessions (Supplementary Data 7,9,10). This mutation was nearly twice as common in post-treatment populations (8.43% of accessions; *n* = 15 miracidia from 10 individuals) compared to pre-treatment populations (4.40% of accessions; *n* = 15 miracidia from 13 individuals). A further 24.9% of accessions had a single copy of this marker. The second and third markers were 150 kb deletions adjacent to *Smp_246790* (∼1.22 Mb) and *Smp_345310* (∼3.18-3.33 Mb), respectively. Structural variant genotyping did not identify the former variant in our accessions, but we did find a series of long homozygous deletions (69.9-215.0 kb) located between 3.02-3.36 Mb in 43.3% of genotyped accessions (*n* = 170 of 393 genotyped accessions; Supplementary Data 11, Supplementary Fig. 10). However, the 150 kb deletion was not enriched in post-treatment populations, and neither the deletion nor p.I1020l clustered phylogenetically (Supplementary Fig. 9).

### Host infrapopulations reveal the extent of genetic relationships and evidence of treatment failure

Sampling and analysis of the parasite population from within a single host (defined as the ‘infrapopulation’) across specific time points has the potential to identify instances of treatment failure, which may be indicative of praziquantel resistance. However, these infrapopulations are thought to be highly heterogeneous (*25*, *47*, *48*), and previous genomic surveys of *S. mansoni* have only included small numbers of parasites from the same individuals (max *n* = 11) (*39*, *41*, *49*, *50*) reducing the likelihood of identifying related parasites. To provide greater resolution of infrapopulation structure and evaluate the impact of praziquantel treatment, we analysed pairwise kinship between 164 sequenced miracidia sampled pre- or post-praziquantel treatment from a total of three donors: Bb1 (*n =* 47 pre-treatment, *n =* 42 post-treatment), Bu3 (*n =* 33 pre-treatment, *n =* 19 post-treatment) and Bu1 (*n =* 2 pre-treatment, *n =* 21 post-treatment) and reanalysed the 174 accessions from the Vianney et al. (*41*) survey (98 donors, *n* = 123 pre-treatment, *n* = 51 post-treatment).

We identified 49 first-degree relationships, 13 second-degree relationships and 34 third-degree relationships (Fig. 4A; Supplementary Data 12). Forty-seven of the 49 first-degree relationships were between accessions sampled from the same donor; the remaining two were between donors and have previously been identified as potentially mislabelled accessions (*41*). Estimates of the coefficient of inbreeding (*f*) using the condensed identity coefficients also revealed low levels of inbred relatedness (median *f* = 0.000227; range *f* = 1×10^-6^ - 0.35) in 18.8% of parents (Supplementary Data 13) (*51*). Within infrapopulations of donors Bb1, Bu1 and Bu3, we identified three first-degree relationships between pre-treatment accessions and 23 between post-treatment accessions. Most of these were found in infrapopulations from Bb1, represented by three independent clusters of four accessions (Fig. 4B,C). We also identified one first-degree and four second-degree relationships spanning treatment arms, suggesting bi- and uni-parental survival, respectively. Both of the first-degree relatives were also homozygous for the *Sm.*TRPM_PZQ_-2723187C marker; however, none of the second-degree relatives had a homozygous or heterozygous copy. Additionally, we did not identify any homo- or heterozygous variants of p.R1843Q in any of the first- or second-degree relatives spanning treatment. We also found no evidence of reduced nucleotide diversity in post-treatment populations (Fig. 4C) and low genetic differentiation between pre- and post-treatment populations (range mean *F*_ST_ = 1.24×10^-3^ - 1.75×10^-3^, Supplementary Fig. 11).

**Fig. 4:**
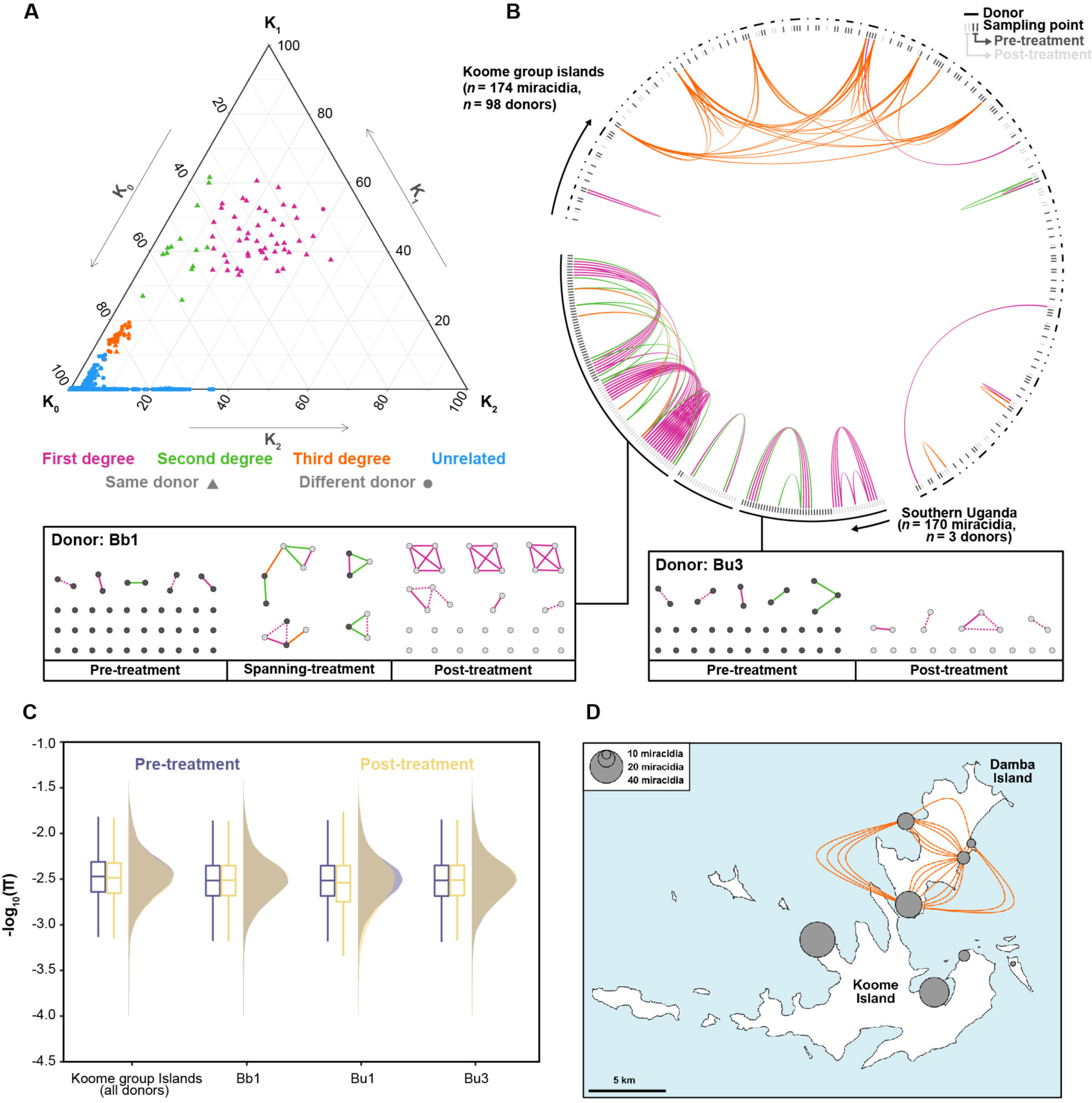
Relatedness between *Schistosoma mansoni* accessions from endemic regions. **A.** Ternary plots representing pairwise relationships between accessions using the three relatedness coefficients *K*_0_, *K*_1_, and *K*_2_, representing the probabilities that at a given locus, the two accessions shared zero, one or two-allele identity-by-descent, respectively. Each point represents a pairwise relationship between two accessions, showing whether each accession was from the same (triangles) or different (circles) donor. Points are coloured by the inferred relationship: first-degree (pink), second-degree (green), third-degree (orange) or unrelated (blue). Only accessions from three Southern Ugandan donors and all Koome Island donors are shown. **B.** Inferred relationships within and between donors. The outer lines represent each donor; within each line, individual accessions are shown coloured by whether they were sampled before (dark grey) or after (light grey) treatment with praziquantel. Coloured lines represent the same pairwise relationships as a). For donors Bb1 and Bu3, all relationships are shown below the circular plot and shown in boxes where they are grouped into pre-treatment samples, post-treatment samples, and pre- and post-treatment samples (spanning treatment). Within the boxes, dotted lines represent relationships inferred by NgsRelate but were found to have a lower degree or no-relation by Sequoia. **C.** The effect of praziquantel treatment on nucleotide diversity (π) was calculated for each donor infrapopulation (Bb1, Bu1 and Bu3) and all Koome group island samples. Nucleotide diversity was calculated in 5 kb non-overlapping windows across each autosome for each pre-treatment (purple) and post-treatment (yellow) population (including related samples). For all boxplots, the central line indicates the median, and the top and bottom edges of the box indicate the 25th and 75th percentiles, respectively. The maximum whisker lengths are specified as 1.5 times the interquartile range. **D.** Identification of a circulating lineage of *S. mansoni* on Damba Island (the northernmost island of the Koome group islands). We identified a cluster of nine accessions, all with third-degree identical by descent relationships, which clustered phylogenetically with accessions from Lake Albert.

Examination of third-degree relationships within the Vianney and colleagues (*41*) dataset identified a cluster of nine related accessions, all localised to Damba Island and found in all four of the sampled villages (Fig. 4B,D). Phylogenetic analysis showed all nine formed a distinct clade with six accessions from Lake Albert (Supplementary Fig. 12), suggesting that these are derived from a recently imported lineage from Lake Albert, recently dispersed across Damba Island.

## DISCUSSION

As efforts to interrupt transmission and eliminate schistosomiasis as a public health problem intensify over the next decade, the escalating use of MDA is expected to result in increased selective pressures, altered transmission dynamics and widespread reductions in schistosome populations (*1*, *9*). The consequences of such large-scale changes to schistosome populations are unclear but provide a strong incentive for genomic surveillance to detect and monitor such changes. In this study, we have, to our knowledge, assembled the most comprehensive collection of *Schistosoma* whole-genome sequencing data to date, including samples from across eight countries and two major foci of infection.

Praziquantel resistance represents a credible threat to schistosomiasis control if it were to establish in endemic populations. Despite over 50 years of use, TRPM_PZQ_ has only recently been identified as a direct target of praziquantel and a genetic determinant of reduced praziquantel susceptibility (*36*, *37*). We identified extensive, low-frequency variation within this gene and used this variation database to inform the targeted mutagenesis of 12 residues, focusing on mutations at potential key residues. We identified four mutations in our population genetic analyses that reduced or ablated channel responsiveness to praziquantel functional genetic *in vitro* assays, and while only one (p.R1843Q) was present in more than a single accession, all represent standing variation for praziquantel resistance in endemic populations. Both p.R1843Q and p.I1020l (the latter a marker for praziquantel resistance in passaged laboratory lines (*37*)) variants were present in multiple accessions, indicating potential variability in praziquantel efficacy in these populations. Both mutations were also more prevalent in post-treatment parasite populations than their pre-treatment counterparts. However, our analyses and others (*23*, *25*, *39*) have shown that post-treatment populations do not typically represent subpopulations of pre-treatment parasites, which may result from reinfection or under-sampling. It is possible that the increased prevalence of these mutations is unrelated to praziquantel administration. It is clear, nevertheless, that despite the asymmetrical per-host sampling of this study, these variants remain widespread in parasite populations sampled from a wide range of hosts, showing that they are prevalent across Lake Victoria. Further work should also be undertaken to determine to what extent the mutations we identified impact the *in vivo* efficacy of praziquantel and parasite fitness. Profiling these specific SNPs in the context of other changes within TRPM_PZQ_ in the same samples, reflecting broader allelic variation, is also necessary.

Our analyses have demonstrated how genomic surveillance of parasites can help identify variants contributing to reduced praziquantel efficacy in endemic populations and provide structural and functional insights into TRPM_PZQ_ that will permit a better understanding of praziquantel efficacy in schistosomes. While we only characterised a small proportion of the total variation we observed, our results survey genetic variation within endemic regions where praziquantel has seen extensive use. These data provide candidate variants for functional investigations and serve as a database of variation for retrospective surveys once further resistance-conferring mutations are identified. However, it is notable that a large proportion of our dataset was derived from donors living in regions of long-term praziquantel administration. Targeted mutagenesis identified four mutations in endemic populations that reduced channel sensitivity to praziquantel, indicating standing variation for resistance. Whilst we did identify both standing variation for future resistance evolution and instances of treatment failures here, the apparent lack of obvious high-frequency resistance-conferring mutations is encouraging as these populations have been under long-term MDA pressure. The variable coverage of MDA may have allowed substantial refugia populations to exist in untreated humans, snails and animal reservoirs. Refugia would reduce the selection pressure for resistance, an approach that has been deliberately employed in veterinary systems (*52–54*). Alternatively, reduced praziquantel susceptibility might be linked to regulatory changes in expression or RNA splicing (*37*), neither of which would be identified by our analyses. Considering how recently *Sm.*TRPM_PZQ_ was recognised as a mediator of praziquantel sensitivity, future functional investigations will likely reveal additional relevant variants.

Despite involving a small number of donors, our analyses of host infrapopulation dynamics represent the most thorough genomic characterisation of a host infrapopulation of any parasitic helminth. Our analyses found that host infrapopulations are exceptionally diverse, consistent with both autopsy studies and genetic surveys, which have found that individual worm burdens range from one up to hundreds or thousands of worm pairs per individual in low and high-endemicity regions, respectively (*55*, *56*). Despite this, we did find evidence of sibling relationships within both treatment arms, with indications of a higher degree of post-treatment relatedness. We also found evidence of post-treatment parental survival, suggesting either ineffective treatment or higher praziquantel tolerance within these populations. However, the limited number of these accessions limited any formal analyses of the possible genetic basis of reduced praziquantel efficacy. Consistent with previous genetic studies, these only represented a small proportion of the overall post-treatment populations (*25*), suggesting that most post-treatment samples represent either infection with additional parasites and/or parasites missed in pre-treatment sampling.

To conclude, we have characterised endemic *S. mansoni* populations at multiple spatial scales ranging from comparisons across major foci of infection to individual parasite infrapopulations. Our variation analysis within a candidate praziquantel resistance loci enabled us to identify multiple novel resistance-conferring mutations. These remain at low frequencies in studied populations but demonstrate standing variation from praziquantel resistance in endemic populations undergoing MDA. Our analyses also provide a resource for retrospective or confirmatory support of functional analyses of this recently identified locus. Finally, our extensive sequencing of host infrapopulations provided an in-depth genomic characterisation of parasite diversity and how this is impacted by praziquantel treatment. Overall, our study summarises *S. mansoni* genomic diversity and represents a resource to assess the efficacy of current interventions and guide future treatment strategies.

## MATERIALS AND METHODS

The data analysed here incorporated whole-genome accessions from published datasets from eight countries and 205 new *Schistosoma* sample data (Fig. 1A; Table 1; Supplementary Data 1). The sections below describe the origin, collection, and ethical approval of samples, focusing only on the parasite samples sequenced for this study and not previously published accessions.

### Collection and ethical approval for the sampling of Ugandan miracidia

The collection of all Ugandan miracidial samples was undertaken as part of the monitoring, evaluation and disease control activities conducted by the Vector Control Division of the Ministry of Health (Uganda), the Schistosomiasis Control Initiative and Imperial College London. All methods and data collection were approved by the Uganda National Council for Science and Technology (Memorandum of Understanding: sections 1.4, 1.5, 1.6) and the Imperial College Research Ethics Committee (EC NO: 03.36. R&D No: 03/SB/033E). The Head of the Vector Control Division informed local district officials, and the headteachers of each school were informed about the study and requested to provide informed consent to allow sampling to be performed within the school. Parents of the children were informed of the study through school meetings, where they were provided with detailed information regarding the purpose of the study and technical staff were present to answer questions. Parents were requested to provide informed consent for their children to participate in the study, and in addition, any children aged ten or older were asked to give informed consent after receiving complete information about the study. Participation was voluntary, access to treatment was not dependent on participation in the study, and children could withdraw or be withdrawn from the study at any time.

Stool samples were collected from each child, and duplicate Kato-Katz thick smears were conducted (*57*). A Pitchford–Visser funnel was used to wash and filter parasite eggs from the remaining stool sample, and the filtrate was stored overnight (*58*). Miracidia were hatched the following day and were transferred into two sequential dishes of nuclease-free water to dilute bacterial contaminants before being individually fixed onto Whatman FTA-indicating classic cards (*59*, *60*). Between 1-3 days following testing, children with evidence of parasitic infection were treated with praziquantel (40 mg/kg) for schistosomiasis and albendazole (400 mg) for soil-transmitted helminths. Children were retested 25-27 days following treatment, and in cases of incomplete *S. mansoni* clearance, miracidia were sampled again, and treatment was re-administered.

### Collection and ethical approval for the sampling of Ugandan *S. mansoni* cercariae

The FTA-preserved Ugandan *S. mansoni* cercariae were collected as part of snail collection surveys conducted between 2007 and 2010 as part of the EU-CONTRAST programme, a consortium of European and African researchers (CONTRAST EU/INCO.Dev contract No. 032203) in association with the Schistosomiasis Control Initiative (*61*) and provided via SCAN. The Ethical Review Board of the Uganda National Council of Science & Technology approved the surveys.

### Collection and ethical approval for the sampling of Tanzanian *S. mansoni* cercariae

Tanzanian *S. mansoni* cercariae were originally collected as part of snail collection surveys undertaken in the Mwanza region and Geita regions of Northern Tanzania between January 2012 and December 2015 (*62*) as part of the Schistosomiasis Consortium for Operational Research and Evaluation (SCORE) snail project (*63*) and archived within the Schistosomiasis Collection at the Natural History Museum (SCAN) (Emery et al., 2007). Survey sites were picked based on their proximity (within 5-15 metres) to schools and were identified based on local information about water activities (bathing, fishing, water collection) (*63*). *Biomphalaria* spp. snails were collected by scooping using handheld metal sieve scoops or dredging using a metal dredge dropped from a boat and dragged 10 m back to shore. Scooped/dredged snails were hand-collected with forceps and placed into collection jars. All snails were placed in 24-well ELISA plates under direct light for at least four hours to induce shedding, and cercariae from each shedding snail were individually collected in 3 μL of water using a Gilson pipette and fixed on Whatman FTA cards (*59*, *60*). COX-1 identification of *S. mansoni* was performed as described in Gouvras *et al.* (*62*).

### Origin of the Senegalese *S. mansoni* laboratory (adult worm) sample

Both *S. manson*i Senegalese isolates (FS0001 and FS0002) originated from approximately 30 *Biomphalaria pfeifferi* with patent *S. mansoni* infections found in the Western principal irrigation canal in the Ndiengue District of Richard Toll (*64*). The strain was originally sampled in 1993 and then maintained at the School of Biological Science, University of Wales (Bangor, United Kingdom) and subsequently in the helminGuard laboratory (Süelfeld, Germany) (*65*).

### Origins of the *S. rodhaini* (adult worms and cercariae) samples

The adult *S. rodhaini* adult worm sample (RZ0001) was provided by SCAN from a laboratory-passaged *S. rodhaini* strain originally isolated from infected *Biomphalaria* snails collected in Burundi in 2000. The sample used here represented the 6th passage through laboratory *Biomphalaria glabrata* and *Mus musculus*, under the Home Office project license numbers 70/4687 (before 2003) and 70/5935 (2003–2008) (*59*).

The two *S. rodhaini* cercariae analysed here originated from the Mwanza region of Tanzania. They were collected as part of the SCORE xenomonitoring project and provided by SCAN. Species identification was confirmed by Sanger sequence analysis of a partial region of the *cox1* gene, the ITS1+2 rDNA region, and a partial region of the 18S rDNA region using the methods described in Pennance *et al*., 2020.

### DNA extraction and sequencing

The DNA from individual miracidia and cercariae were isolated from the Whatman FTA cards using methods described by (*66*, *67*). A 2mm Harris micro-punch was used to punch out the FTA disc containing the DNA. Within a well of a 96 PCR plate, individual samples on individual punches were lysed in 30 µL of the following buffer: 30 mM Tris–HCl pH 8.0 (Sigma Aldrich), 0.5% Tween 20 (Sigma Aldrich), 0.5% NP40/IGEPAL CA-630 (Sigma Aldrich) and 1.25 µg/mL of Protease reagent (Qiagen; cat 19155). Punches were incubated at 50°C for 1 h, then heated to 75°C for 30 mins. DNA was extracted from adult worms using the Qiagen MagAttract HMW kit (PN-67653) following the manufacturer’s instructions. The DNA from the two individual Senegalese adult worms was extracted using the Qiagen MagAttract HMW kit (PN-67653) following the manufacturer’s instructions.

Library preparation was performed using a low-input enzymatic fragmentation-based library preparation method (*68*). For each sample, 20 μL of lysate (or extracted DNA) was mixed with 50 μL of TE buffer (Ambion 10 mM Tris-HCL, 1 mM EDTA) and 50 μL of Ampure XP beads followed by a 5 min binding reaction at room temperature. Magnetic bead separation was used to separate genomic DNA, which was then washed twice with 75% ethanol. Beads were resuspended in 26 μL of TE buffer. To perform DNA fragmentation and A-tailing, each sample was immediately mixed with 7 μL of 5x Ultra II FS buffer and 2 μL of Ultra II FS enzyme and incubated on a thermal cycler for 12 min at 37°C followed by 30 min at 65°C. Adaptor ligated libraries were prepared by adding 30 μL of ligation mix, 1 μL ligation enhancer (New England BioLabs), 0.9 μL nuclease-free water (Ambion) and 0.1 μL duplexed adapters to each well followed by incubation for 20 min at 20°C. Libraries were purified and eluted by adding 65 μL of Ampure XP beads and 65 μL of TE buffer. Libraries were amplified by adding 25 μL KAPA HiFi HotStart ReadyMix (KAPA Biosystems) and 1 μL PE1.0 primer to 21.5 μL of the library. Each sample was thermal-cycled as follows: 98 °C for 5 min, then 12 cycles of 98°C for 30 sec, 65°C for 30 sec, 72°C for 1 min and finally 72°C for 5 min. Ampure beads were used to purify amplified libraries using a 0.7:1 volumetric ratio of beads to library. Each library was then eluted into 25 μL of nuclease-free water. Library concentrations were adjusted to 2.4 nM and pooled, followed by sequencing on the Illumina NovaSeq 6000 using 150 bp paired-end chemistry.

### Variant discovery and annotation

In addition to the whole-genome sequencing of samples described above, sequence data for 407 *S. mansoni* accessions (*39–41*) and one *S. rodhaini* accession (*40*) were included in this study. Raw sequencing reads from all 682 accessions were trimmed using BBDuk (https://jgi.doe.gov/data-and-tools/bbtools/bb-tools-user-guide/) to remove low-quality bases and adapter sequences. Trimmed sequence reads were aligned to the *S. mansoni* (SM_V9, WormBase ParaSite v16) (*69*) reference genome using BWA mem (v.0.7.17) (*70*). PCR duplicates were marked using PicardTools MarkDuplicates (as part of GATK v.4.2.0.0) (*71*). Variant calling was performed per sample using GATK HaplotypeCaller (v.4.2.0.0) in gVCF mode, retaining both variant and invariant sites. Individual gVCFs were merged using GATK CombineGVCFs, and joint-call cohort genotyping was performed using GATK GenotypeGVCFs. Variant sites with only single-nucleotide polymorphisms (SNPs) were separated from indels and mixed sites (variant sites that had both SNPs and indels called) using GATK SelectVariants. GATK VariantFiltration was used to filter both groups independently. SNPs were retained if they met the following criteria: QD ≥ 2.0, FS ≤ 60.0, MQ ≥ 40.0, MQRankSum ≥ −12.5, ReadPosRankSum ≥ −8.0, SOR ≤ 3.0. Variant sites containing indels or mixed sites were retained if they met the following criteria: QD ≥ 2.0, FS ≤ 200.0, ReadPosRankSum ≥ −20.0, SOR ≤ 10.0.

VCFtools (v.0.1.15) was used to exclude accessions with a high rate of variant site missingness (missing genotype called at >5% of sites) and subsequently used to remove sites where >10% of accessions had a missing genotype (*72*). This formed the primary VCF file used for almost all analyses. For analyses of nucleotide diversity and fixation index (*F*_ST_), we produced a second filtered VCF file. We used VCFtools to filter both variant and invariant sites with >80% missing variants, enforced a minimum mean read depth of 5, a maximum mean read depth of 500, removing variant sites found to be significantly out of Hardy-Weinberg equilibrium (*p* < 0.001), and only retained SNPs. Functional annotation of SNPs and indels in the primary VCF file was performed using SnpEff (v.5.0e) (*73*) with gene annotations (v.9) downloaded from WormBase ParaSite (*74*) V17. Repetitive elements in the *S. mansoni* assembly were annotated using RepeatModeler and RepeatMasker (*75*).

### Depth of coverage

For each sample, the depth of read coverage was calculated in 2 kb windows along each chromosome using bedtools coverage (v.2.30.0) (*76*).

### Sample relatedness

We estimated pairwise relatedness between accessions from the same endemic regions. We subsampled to only autosomal SNPs and ran NgsRelate (*77*) using default parameters. As an additional line of evidence, we first used PLINK (v.2.0) (*78*) to exclude SNPs found to be in strong linkage disequilibrium. The genome was then scanned in sliding windows of 50 SNPs, increasing in steps of 10 SNPs and SNPs within windows with squared correlation coefficients >0.2 were removed. Using this filtered subset of SNPs, we performed pedigree reconstruction using Sequoia (*79*). Pairwise relationships were first classified based on the coefficient of kinship score (θ) calculated by NgsRelate. NgsRelate was first run on all 570 *S. mansoni* samples to identify relationships between datasets or populations. After confirming these relationships did not occur, NgsRelate was rerun on individual populations from Southern Uganda, Northern Tanzania and Koome Islands. Additionally, samples from Berger et al. (*39*) were excluded from these analyses due to both highly elevated inbreeding coefficients in this dataset. Pairwise relationships with θ > 0.354 were classified as monozygotic twins, 0.354 > θ ≥ 0.177 were classified as first-degree relatives, 0.177 > θ ≥ 0.0884 were classified as second-degree relatives and 0.0884 > θ ≥ 0.0442 were classified as third-degree relatives. In addition, first-, second-, and third-degree relationships were only considered valid if the maximum likelihood estimate of sharing 1 IBD allele (K_1_) was greater than 0.05. Instances where both NgsRelate and Sequoia identified the same first-degree relationships were designated ‘high-confidence’ relationships. GGtern was used to plot all three maximum likelihood estimates of sharing 0, 1 or 2 IBD alleles (K_0_, K_1_, and K_2_), and Circos (*80*) and Gephi (*81*) were used to visualise pairwise relationships between different donors.

### Population genomic structure and diversity

We removed variants found at minor allele frequencies < 0.05 and excluded all variants found on the Z chromosome, W chromosome and mitochondrial genome. We then removed all variants found within repetitive regions identified using RepeatMasker and finally removed variants found to be in strong linkage disequilibrium, as above. Principal component analysis was performed with the remaining 188,923 autosomal SNPs using PLINK. Admixture analyses were performed using ADMIXTURE (*82*) with K values (number of hypothetical ancestral populations) ranging from 1 to 20, 10-fold cross-validation and standard error estimation with 250 bootstraps. The lowest cross-validation error (CV) value was found for K = 5 (Supplementary Fig. 13).

We used publicly-available scripts to convert all 188,923 autosomal SNPs into Phylip format (https://github.com/edgardomortiz/vcf2phylip/vcf2phylip.py) and remove invariant sites (https://github.com/btmartin721/raxml_ascbias/ascbias.py).

Phylogenomic inference was performed using IQ-TREE (*83*), using the best-fit substitution model with ascertainment bias correction selected by ModelFinder (GTR+F+ASC+R10) and 1000 ultrafast bootstraps. The resulting phylogeny was visualised using ggtree (*84*).

pixy (v.1.2.3.beta1) (*85*) was used to calculate autosomal nucleotide diversity (π) and the fixation index (*F*_ST_) in 5 kb non-overlapping, sliding windows for each population using the secondary (mixed variant and invariant sites) VCF. Negative *F*_ST_ values were corrected to 0 before calculating genome-wide median values. Watterson’s estimator (Θ) was calculated using Scikit-allel (*86*). Effective population size (N_e_) was using a per-generation mutation rate (μ) of 8.1×10^−9^(*40*) and the following equation:

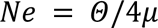

The coefficient of inbreeding (F) was calculated using VCFtools to assess per-sample homozygosity. Comparisons of samples within and between study sites can be found in Supplementary Fig. 14A-C.

### The impact of praziquantel treatment on population genomic diversity

pixy (v.1.2.3.beta1) was used to calculate autosomal nucleotide diversity (π) in 5 kb non-overlapping, sliding windows for each pre- and post-treatment population using the secondary (mixed variant and invariant sites) VCF and including related accessions (*85*).

### Structural variant genotyping and annotation

Structural variants for each sample were detected using LUMPY (v.0.2.13) and genotyped using SVTyper (v.0.7.0) as implemented in the Smoove (v.0.2.7) pipeline (https://github.com/brentp/smoove) (*87*, *88*). Using this pipeline, we first performed single-sample calling using the BWA-aligned reads to the *S. mansoni* reference genome. This was followed by merging called variants and sample-wise re-genotyping across all 570 *S. mansoni* accessions. Structural variants intersecting coding regions were annotated using the reference GFF file. We then filtered variants based on the Smoove author’s recommendations, where heterozygous calls with MSHQ>=3, deletions with DHFFC>=0.7 and duplications with DHFFC<=1.25 were excluded. Due to inconsistent coverage and whole-genome amplification of the original libraries, all accessions from Berger et al. (*39*) were excluded from structural variant analyses after genotyping.

### Inference of demographic history

We ran SMC++ (v.1.15.2) (*89*) on each autosome using a per-generation mutation rate of 8.1×10^−9^ and a generation time of 85 days. For Southern Ugandan, Koome Island, Northern Tanzanian, and Eastern Ugandan populations, we randomly subset them down to *n* = 12 unrelated accessions, providing replicates where populations had more than 24 accessions. For all other populations, only single accessions were analysed.

### Screening for recent admixture

Analysis of passaged laboratory strains also identified a single probable *S. mansoni*-*S.rodhaini* hybrid (RZ0001), which displayed high heterozygosity, intermediate admixture proportions and intermediate phylogenetic positioning, consistent with an early-generation (F_1_ - F_3_) hybrid, this sample was excluded from further analyses (Supplementary Fig. 15; Supplementary Data 2) (*90*, *91*).

### Analysis of candidate praziquantel resistance genes

We inspected SnpEff annotated variants within the candidate praziquantel resistance gene (*Smp_24790*), focusing on isoform 5 (*Smp_246790.5* in the v9 annotation), the only full-length isoform containing the predicted binding site. Frequencies of functionally impactful mutations were calculated using only Lake Victoria accessions. The maximum-likelihood phylogeny produced previously was annotated to highlight accessions containing specifically functionally impactful mutations. We excluded a single indel, p.Met1fs, which was a frameshift variant and a short repeat from the analysis; for completeness, information on this variant is still reported in Supplementary Data 7,9,10.

### Ca^2+^ reporter assays

Ca^2+^ imaging assays were performed using a Fluorescence Imaging Plate Reader (FLIPR^TETRA^, Molecular Devices). HEK293 cells (naïve or transfected with specific TRPM_PZQ_ variants) were seeded (20,000 cells/well) in a black-walled clear-bottomed poly-d-lysine coated 384-well plate (Corning) in DMEM growth media supplemented with 10% FBS. This medium was removed after 24hrs, and the cells were loaded with a fluorescent Ca^2+^ indicator (Fluo-4 NW dye, Invitrogen) by incubation (20 µL per well, 1 h at 37°C) in Hanks’ balanced salt solution (HBSS) assay buffer containing probenecid (2.5 mM) and HEPES (20mM). Dilutions of praziquantel (Sigma) were prepared in this same buffer, but without probenecid and dye, in flat shape 384-well plates (Greiner Bio-one, Germany). The Ca^2+^ reporter assay was performed at room temperature by monitoring fluorescence (raw fluorescence units) before (basal, 20s) and after the addition of praziquantel (an additional 250s). For quantitative analyses, peak fluorescence in each well was normalised to the maximal response of the reference channel sequence and concentration-response curves were plotted using the sigmoidal dose-response function in Origin.

## List of Supplementary Materials

Supplementary Data 1 to 14

Supplementary Figs 1 to 15

## Acknowledgements

We would like to thank all the technicians, nurses and drivers who supported sample collections, including (but not exclusive to) Moses Arinaitwe, Aida Wamboko, Annet Enzaru, Andrina Nankasi, Aaron Atuhaire and Fiddi Rugigana from the Vector Control Division, Ugandan Ministry of Health, Uganda, and John Igogote, Honest Nagai, and Revocatus Alphonce from the National Institute for Medical Research, Tanzania. We would also like to thank the teachers from the schools we visited for their assistance and cooperation, the officials in the Health and Education Departments in districts of Mayuge (Uganda), and Alan Fenwick and Wendy Harrison of the Schistosomiasis Control Initiative (renamed Unlimit Health) for their help organising and funding the Ugandan fieldwork. Finally, we thank all the children recruited into this study and their parents and teachers for their cooperation.

## Author contributions

DJB conceived and designed the study with input from JAC, MR, AE, FA, SB, MB, JSM and JPW. TC, NBK, EMT, PHLL and JPW planned and coordinated the collection of samples from Bugoto and Bukagabo Beach schools, which was carried out by TC, PHLL and MA. SCORE Tanzanian samples were collected by AG, BLW, FA, HN, JI, MR, RA, SK, and JPW in partnership with local technicians. CONTRAST samples were collected by CS, JRS, BW, and JPW in partnership with local technicians. JTV, RS and AE provided information on the Koome Island samples. MR, AE and FA curated the SCAN collection and coordinated sample delivery. HH coordinated the isolation and delivery of the two passaged Senegalese laboratory isolates, and NH performed DNA extraction and coordinated sequencing for those samples. For all other samples, DJB extracted and amplified DNA, and planned and coordinated the sequencing. JSM and SKP performed Ca^2+^ reporter assays and modelling analysis. DJB analysed all other data and wrote the paper with input from JSM, SRD and JPW. S-KP, TC, JAC, MR, AE, BLW, PHL, SB and MB provided comments on the text, and all authors viewed and approved the final manuscript.

## Data and materials availability

The sequencing data generated in this study have been deposited in the European Nucleotide Archive (ENA) repository under project accession codes PRJEB42451 and PRJEB29904. Individual sample accessions are listed in Supplementary Data 14. The genome assembly and annotation files are available through ENA (PRJEA36577; GCA_000237925). The code used for data analysis is available at https://github.com/duncanberger/SM_VARIATION.

