## Supplementary Figures for "Extensive transmission and variation in a functional receptor for praziquantel resistance in endemic *Schistosoma mansoni*"

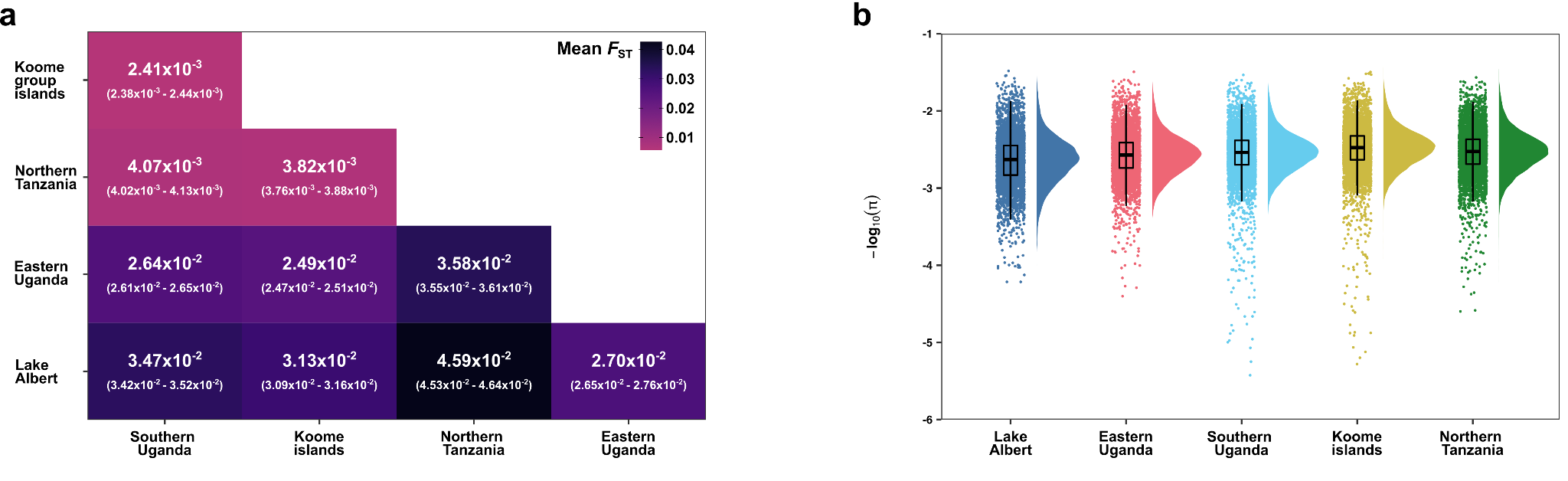

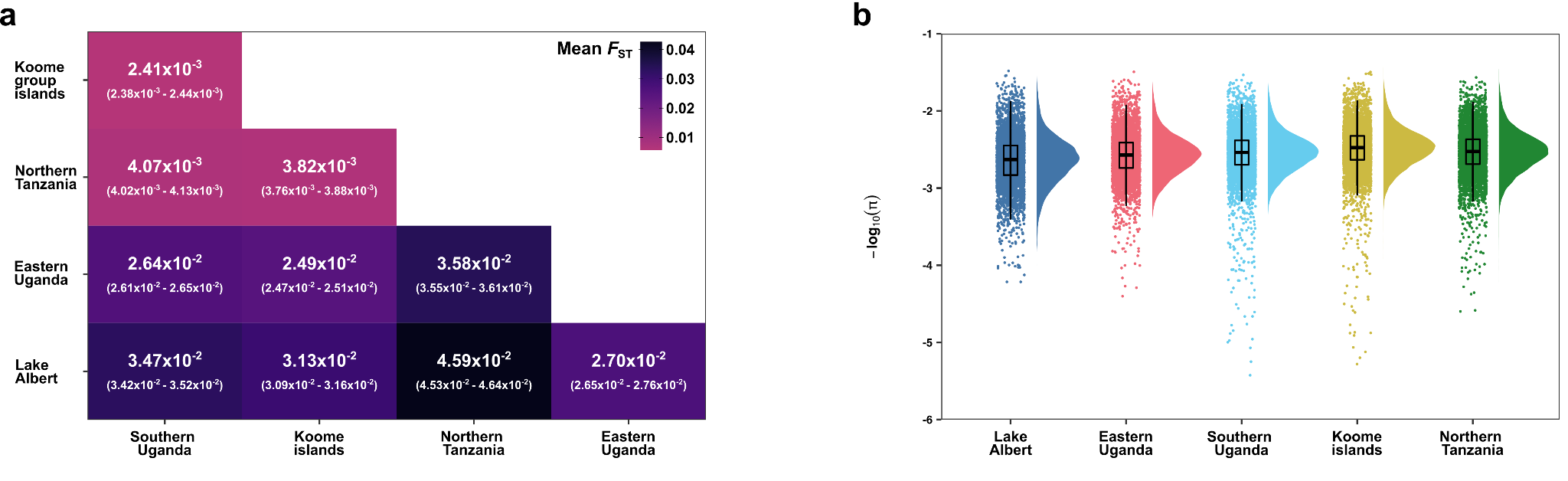

**Supplementary Figure 1. Diversity and differentiation of Lake Victoria *Schistosoma mansoni* populations. a).** Pairwise comparisons of fixation index (*F*_ST_) between each population with greater than two unrelated accessions representing Lake Albert (n = 3 accessions), Eastern Uganda (n = 17), Southern Uganda (n = 287), Koome group islands (n = 159) and Northern Tanzania (n = 30). *F*_ST_ was calculated in calculated using autosomal variants in non-overlapping 5 kb windows. Mean values for each comparison are shown in bold, numbers in parentheses represent the 95% bootstrap confidence intervals around the mean. **b).** Autosomal nucleotide diversity (π) values are calculated as the mean of non-overlapping 5 kb windows for each population described in d). For all boxplots, the central line indicates the median, the top and bottom edges of the box indicate the 25th and 75th percentiles, respectively. The maximum whisker lengths are specified as 1.5 times the interquartile range.

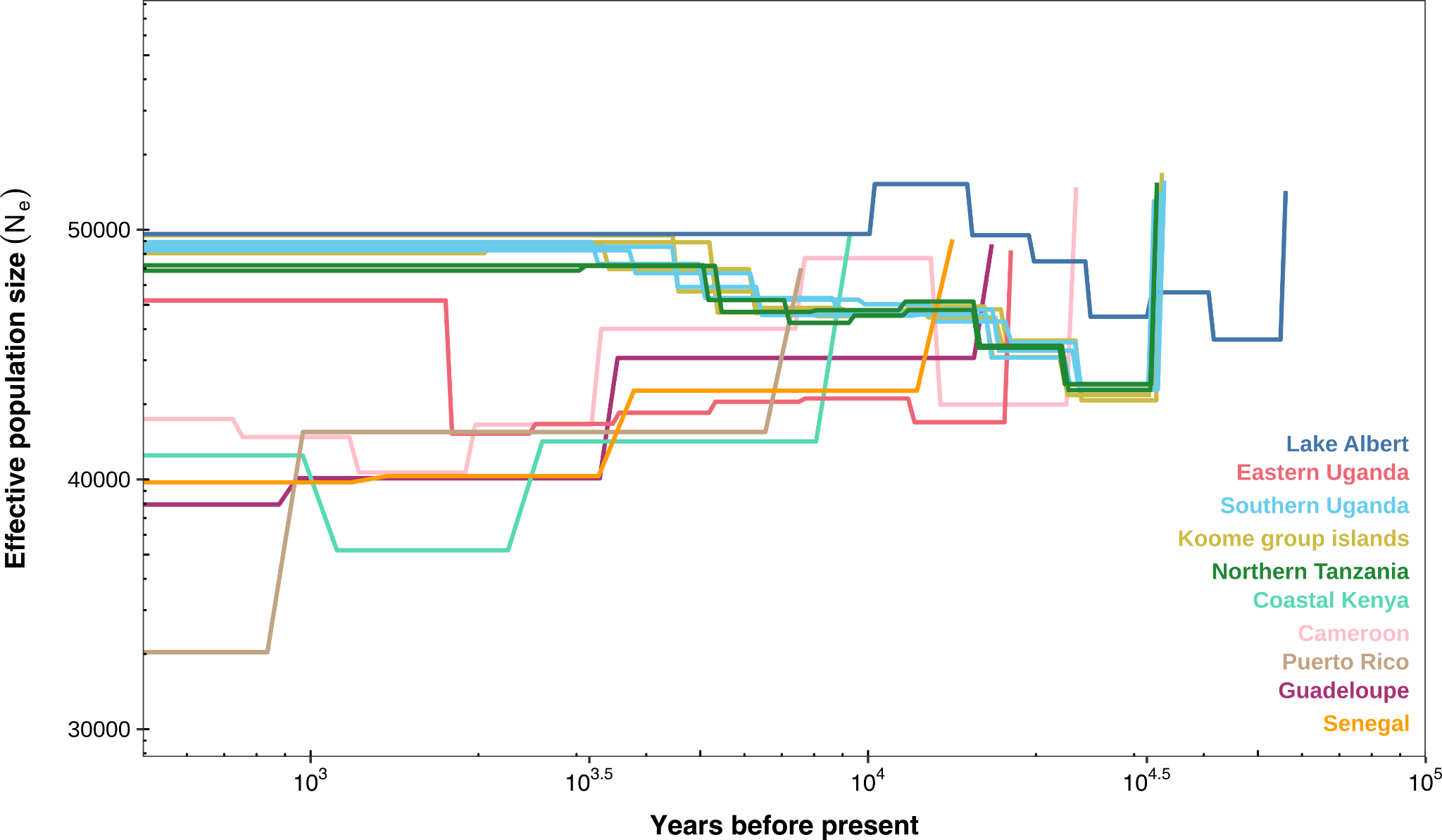

**Supplementary Figure 2. Inference of demographic history using SMC++ for *Schistosoma mansoni* subpopulations**. SMC++ was run on each autosome using a per-generation mutation rate of 8.1x10^-9^ and a generation time of 85 days. We randomly subset populations down to *n* = 12 unrelated accessions for Eastern Ugandan, Southern Ugandan, Koome group island and Northern Tanzanian populations, providing replicates where populations had more than 24 accessions. For all other populations, lines represent single accessions.

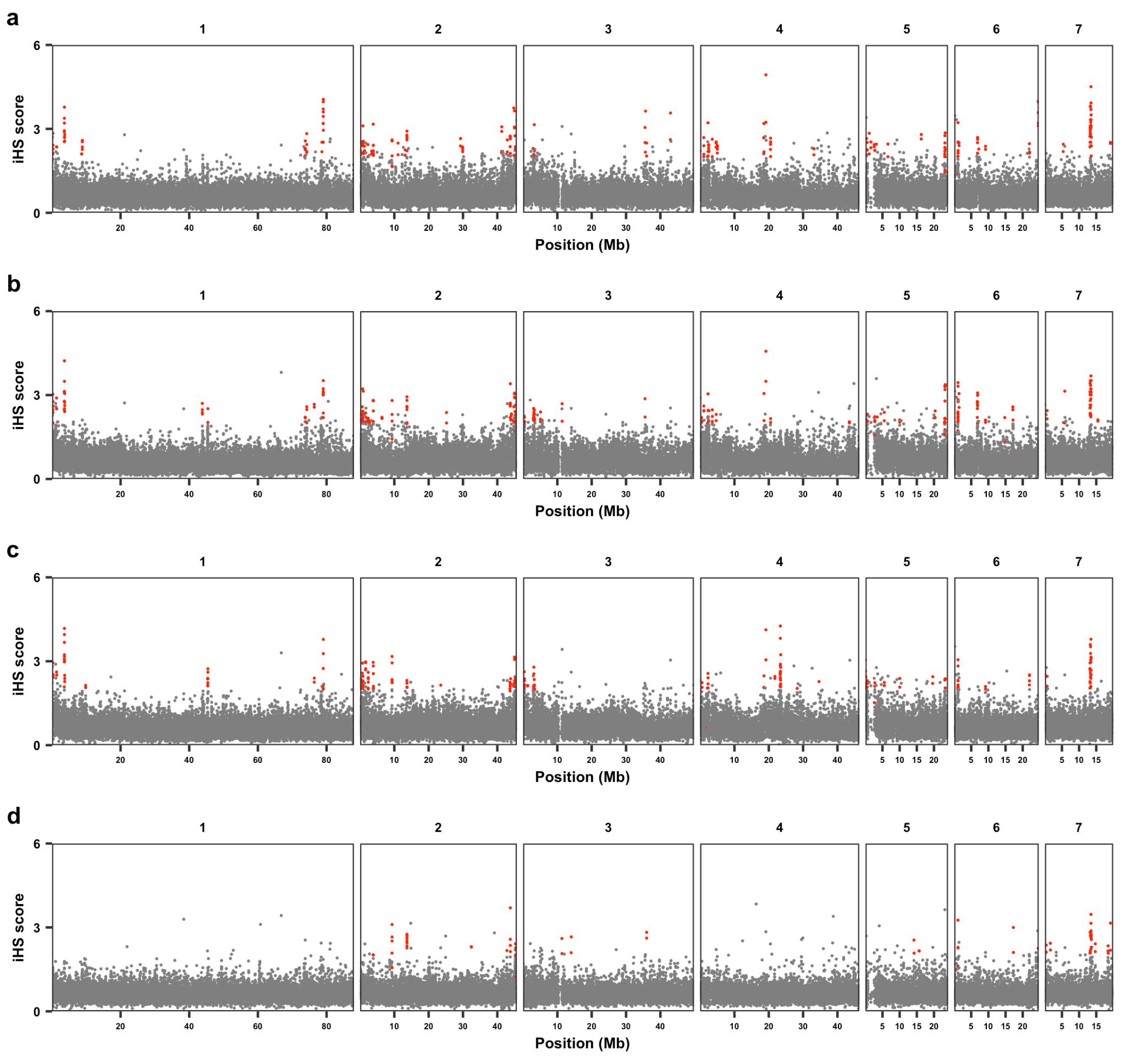

**Supplementary Figure 3. Genome-wide integrated haplotype scores (iHS) were calculated independently using unrelated accessions from four populations.** **a)** Southern Uganda (*n* = 287), **b)** Koome group islands (*n* = 159), **c)** Northern Tanzania (*n* = 30) and **d)** Eastern Uganda (*n* = 17). Points represent median |iHS| values of all variants in 5 kb non-overlapping windows along the seven *S. mansoni* autosomes (grey). Windows with elevated iHS scores (|iHS|>2) within 50 kb of another window were grouped into continuous regions of selection, windows within those regions are shown in red.

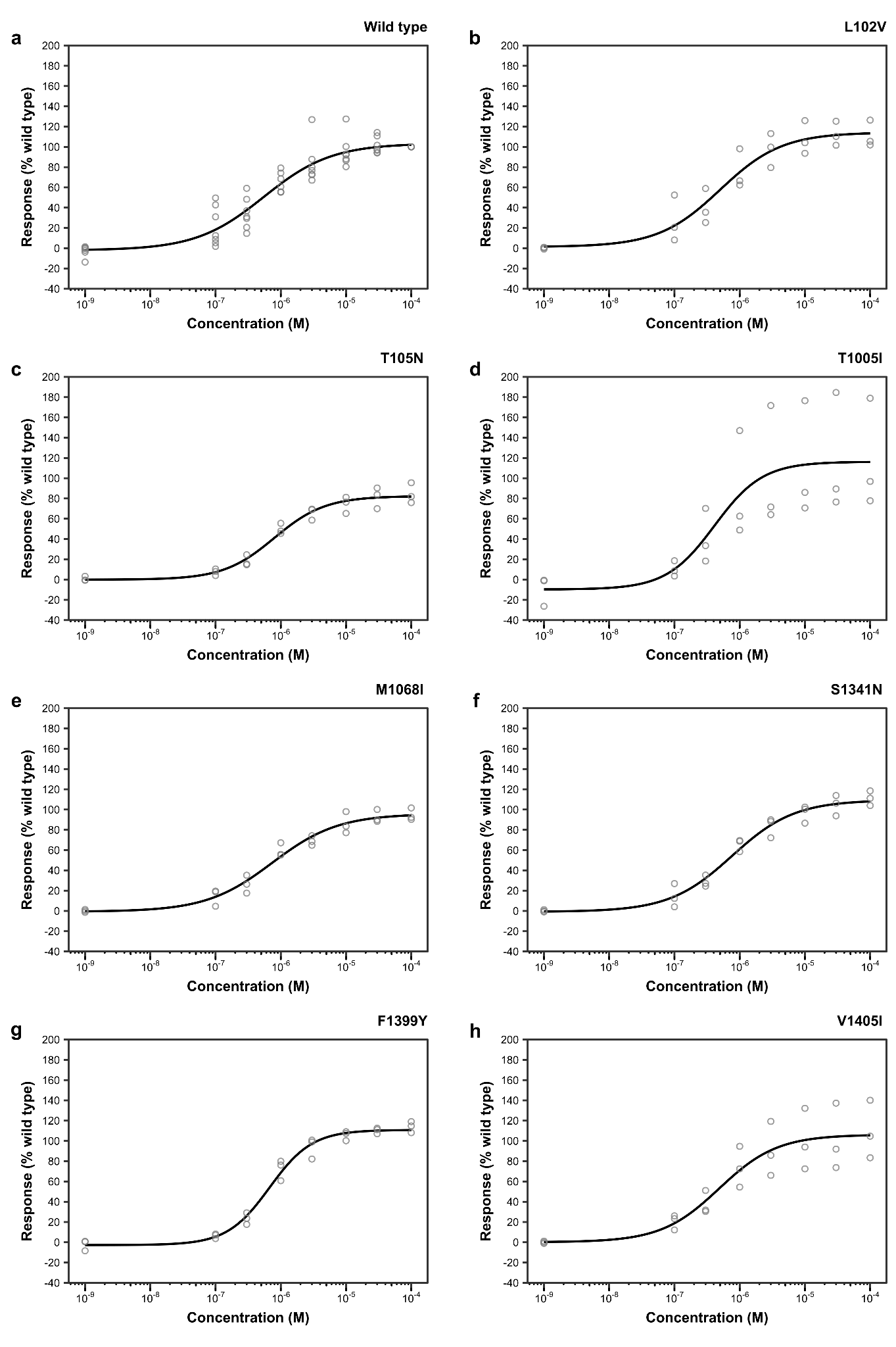

**Supplementary Figure 4. Functional profiling of *Sm*.TRPM_PZQ_ variants.** Concentration-response relationships for the consensus *Sm*.TRPM_PZQ_ sequence compared with seven *Sm*.TRPM_PZQ_ variants. Results represent mean response relative to a) wild type from at least three independent transfections (points). Lines represent the fitted dose-response model using the four-parameter log-logistic function implemented in the drc package.

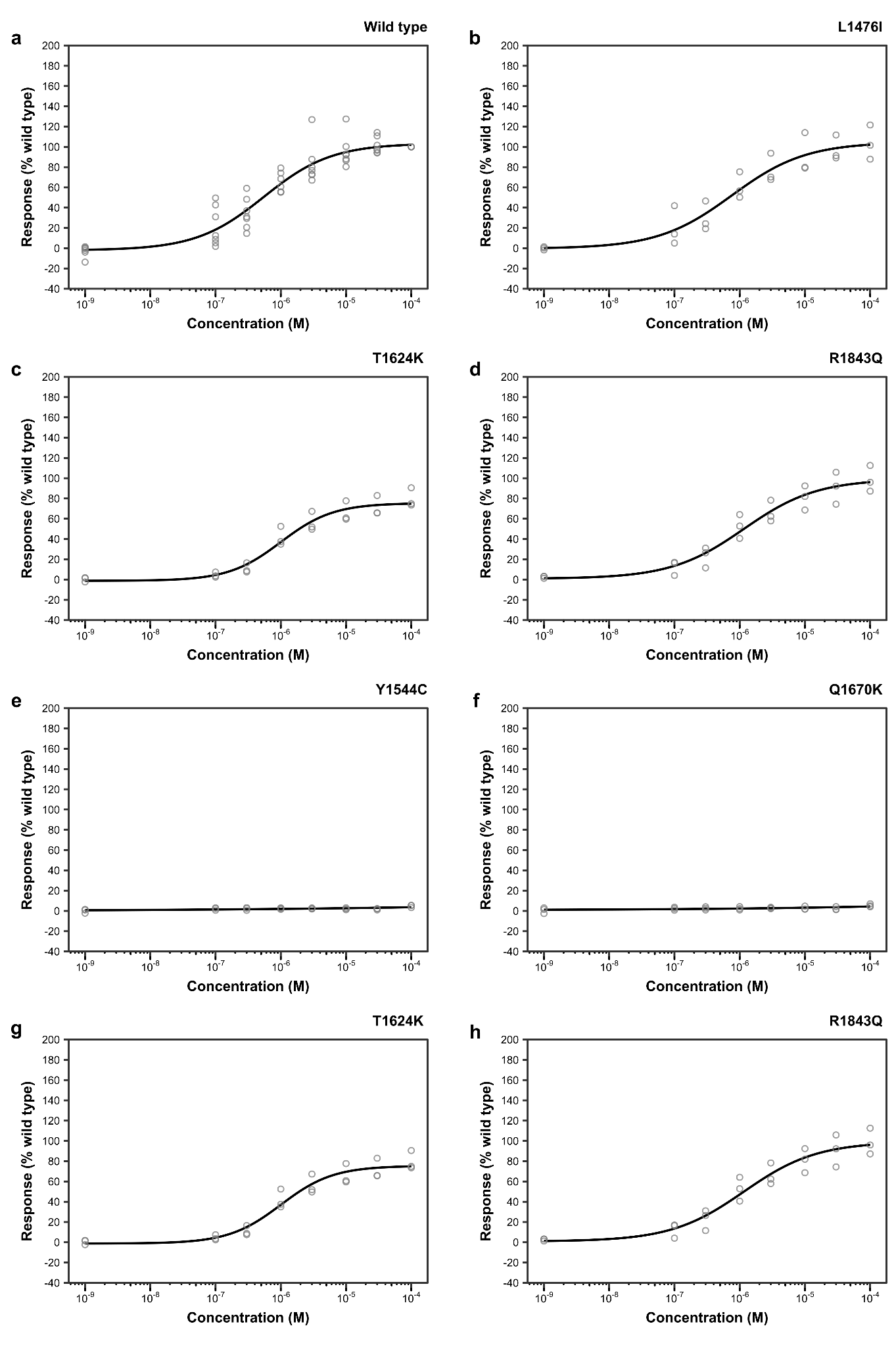

**Supplementary Figure 5. Functional profiling of *Sm*.TRPM_PZQ_ variants.** Concentration-response relationships for the consensus *Sm*.TRPM_PZQ_ sequence compared with six *Sm*.TRPM_PZQ_variants. Results represent the mean response relative to wildtype (Fig. 8a) from at least three independent transfections (points). The lines represent a fitted dose-response model using the four-parameter log-logistic function implemented in the drc package.

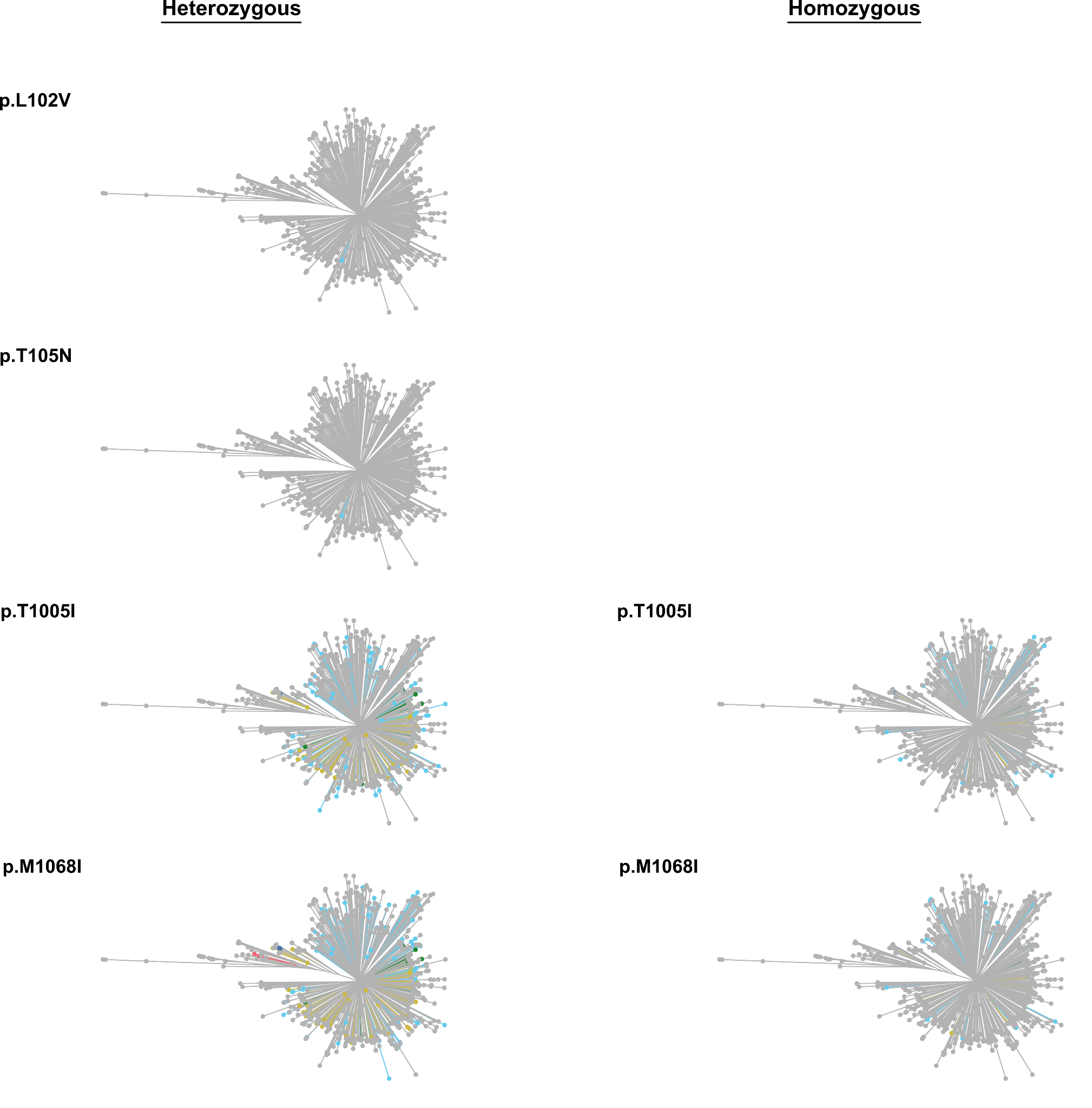

**Supplementary Figure 6. Distribution of individual mutations within *Sm.TRPM_PZQ_* in analysed *Schistosoma* populations.** For each mutation (top left), non-grey lines and points represent accessions which contain the associated variants in either heterozygous or homozygous form. Colours correspond to sampling locations: Puerto Rico (PR; brown), Guadeloupe (GP; orange), Senegal (SN; dark purple), Cameroon (pink), Coastal Kenya (KE; teal), Lake Albert (dark blue), Eastern Uganda (red), Southern Uganda (light blue), Koome group islands (yellow), Northern Tanzania (dark green).

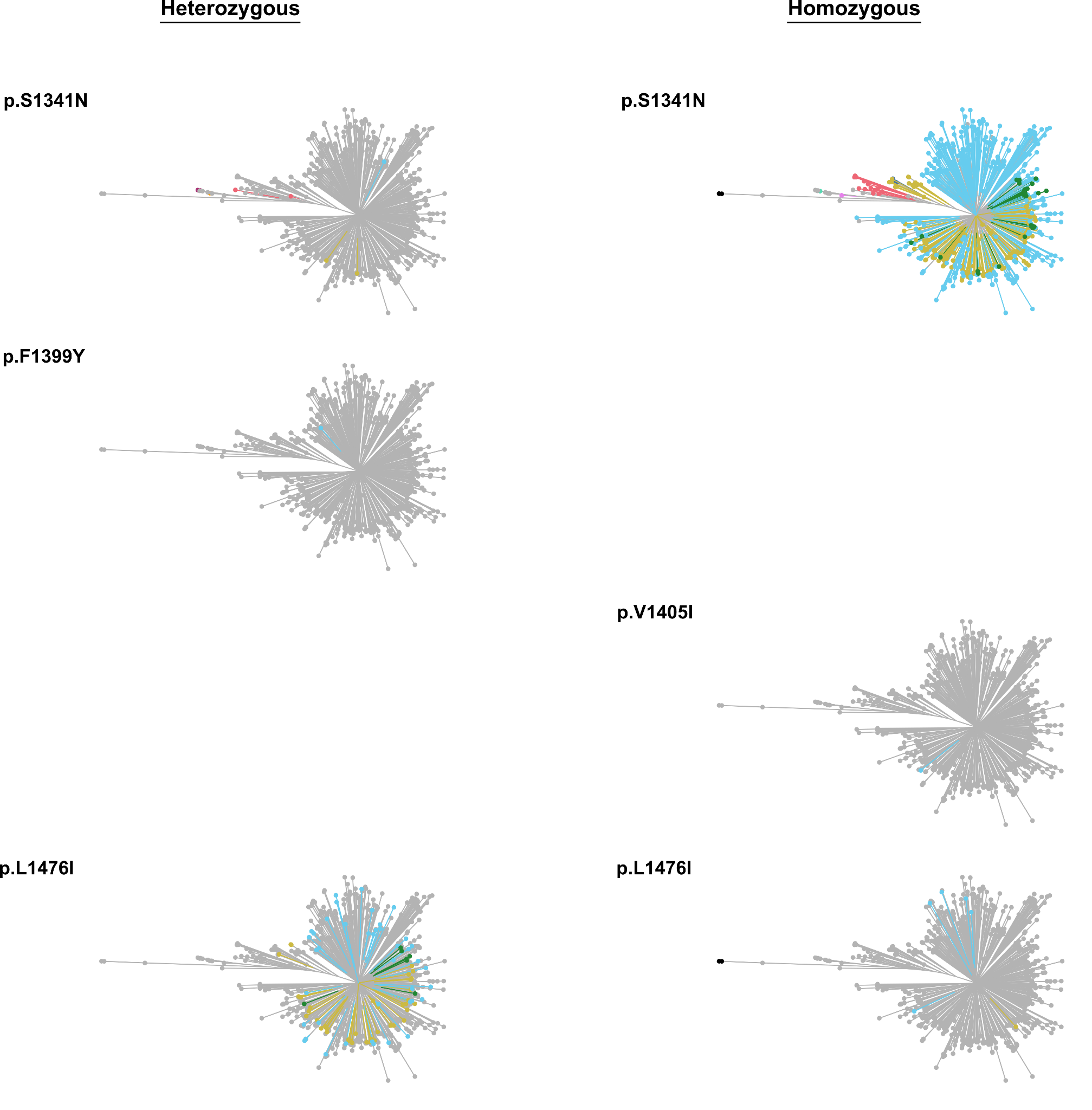

**Supplementary Figure 7. Distribution of individual mutations within *Sm.TRPM_PZQ_* in analysed *Schistosoma* populations.** For each mutation (top left), non-grey lines and points represent accessions which contain the associated variants in either heterozygous or homozygous form. Colours correspond to sampling locations: Puerto Rico (PR; brown), Guadeloupe (GP; orange), Senegal (SN; dark purple), Cameroon (pink), Coastal Kenya (KE; teal), Lake Albert (dark blue), Eastern Uganda (red), Southern Uganda (light blue), Koome group islands (yellow), Northern Tanzania (dark green).

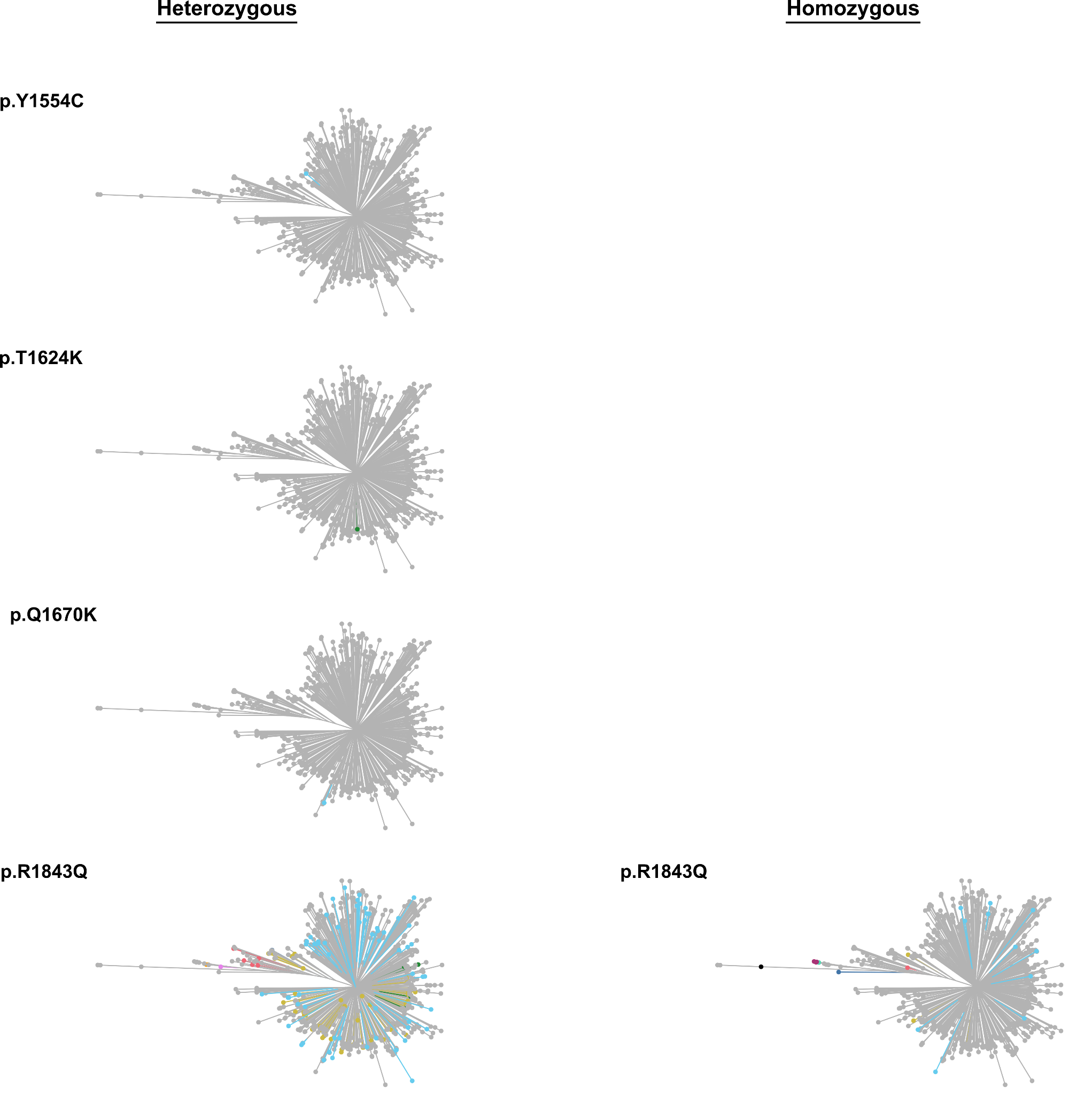

**Supplementary Figure 8. Distribution of individual mutations within *Sm.TRPM_PZQ_* in analysed *Schistosoma* populations.** For each mutation (top left), non-grey lines and points represent accessions which contain the associated variants in either heterozygous or homozygous form. Colours correspond to sampling locations: Puerto Rico (PR; brown), Guadeloupe (GP; orange), Senegal (SN; dark purple), Cameroon (pink), Coastal Kenya (KE; teal), Lake Albert (dark blue), Eastern Uganda (red), Southern Uganda (light blue), Koome group islands (yellow), Northern Tanzania (dark green).

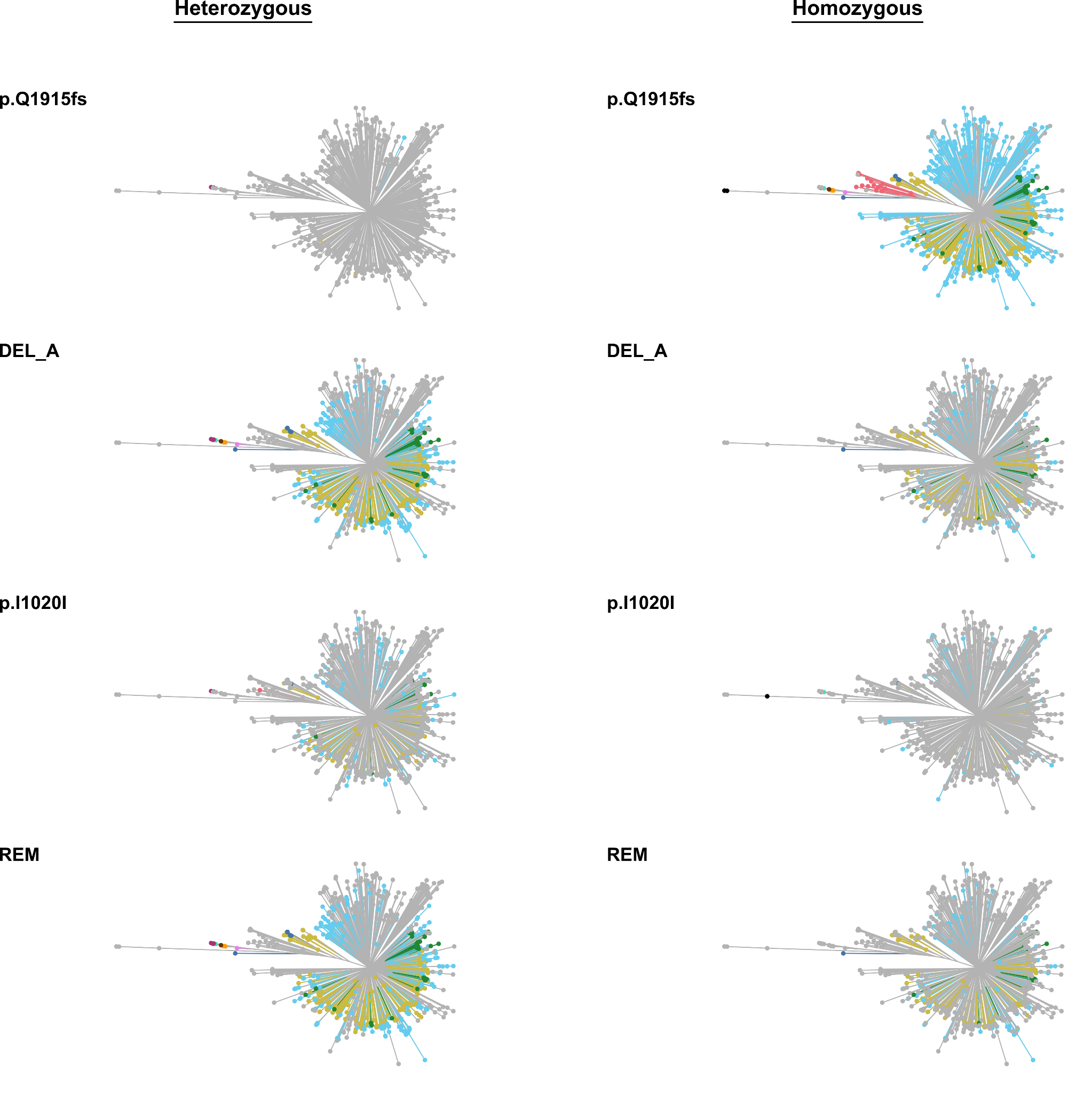

**Supplementary Figure 9. Distribution of individual mutations within or nearby to *Sm.TRPM_PZQ_* in analysed *Schistosoma* populations.** For each mutation or structural variant (top left), non-grey lines and points represent accessions which contain the associated variants in either heterozygous or homozygous form. ‘150 kb deletion’ refers to a series of 69.9-215.0 kb deletions located adjacent to Smp_345310 (~3.18-3.33 Mb on chromosome 3). Colours correspond to sampling locations: Puerto Rico (PR; brown), Guadeloupe (GP; orange), Senegal (SN; dark purple), Cameroon (pink), Coastal Kenya (KE; teal), Lake Albert (dark blue), Eastern Uganda (red), Southern Uganda (light blue), Koome group islands (yellow), Northern Tanzania (dark green).

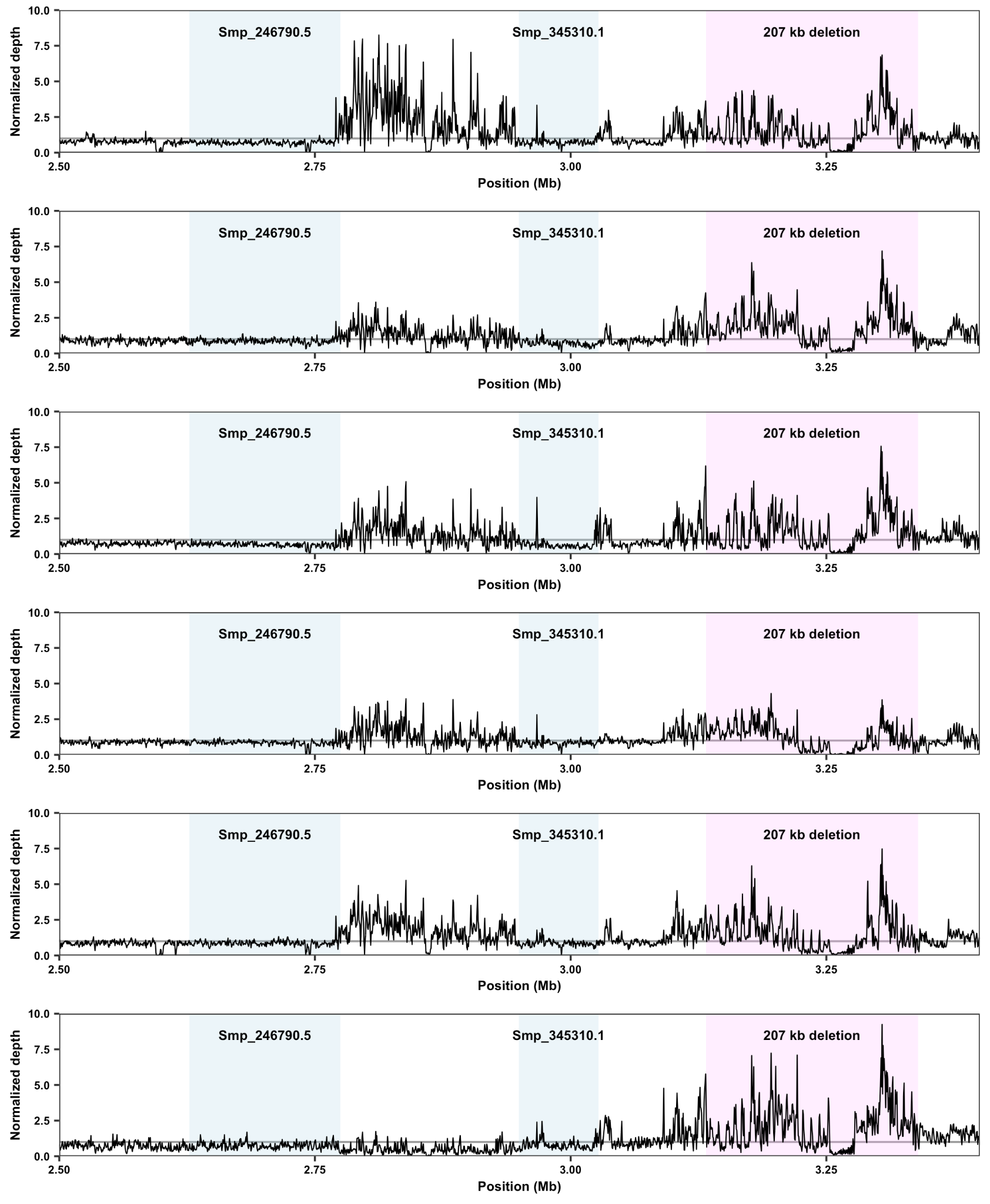

**Supplementary Figure 10.** Depth of coverage of six accessions encompassing two genes implicated in reduced praziquantel susceptibility (Smp_246790.5, Smp_345310.1) and showing genotyped structural variants. Light blue regions indicate the location of each gene and pink regions indicate the range of the genotyped deletion. Black lines indicate the median depth of coverage in 500 bp windows divided by the average autosomal depth of coverage.

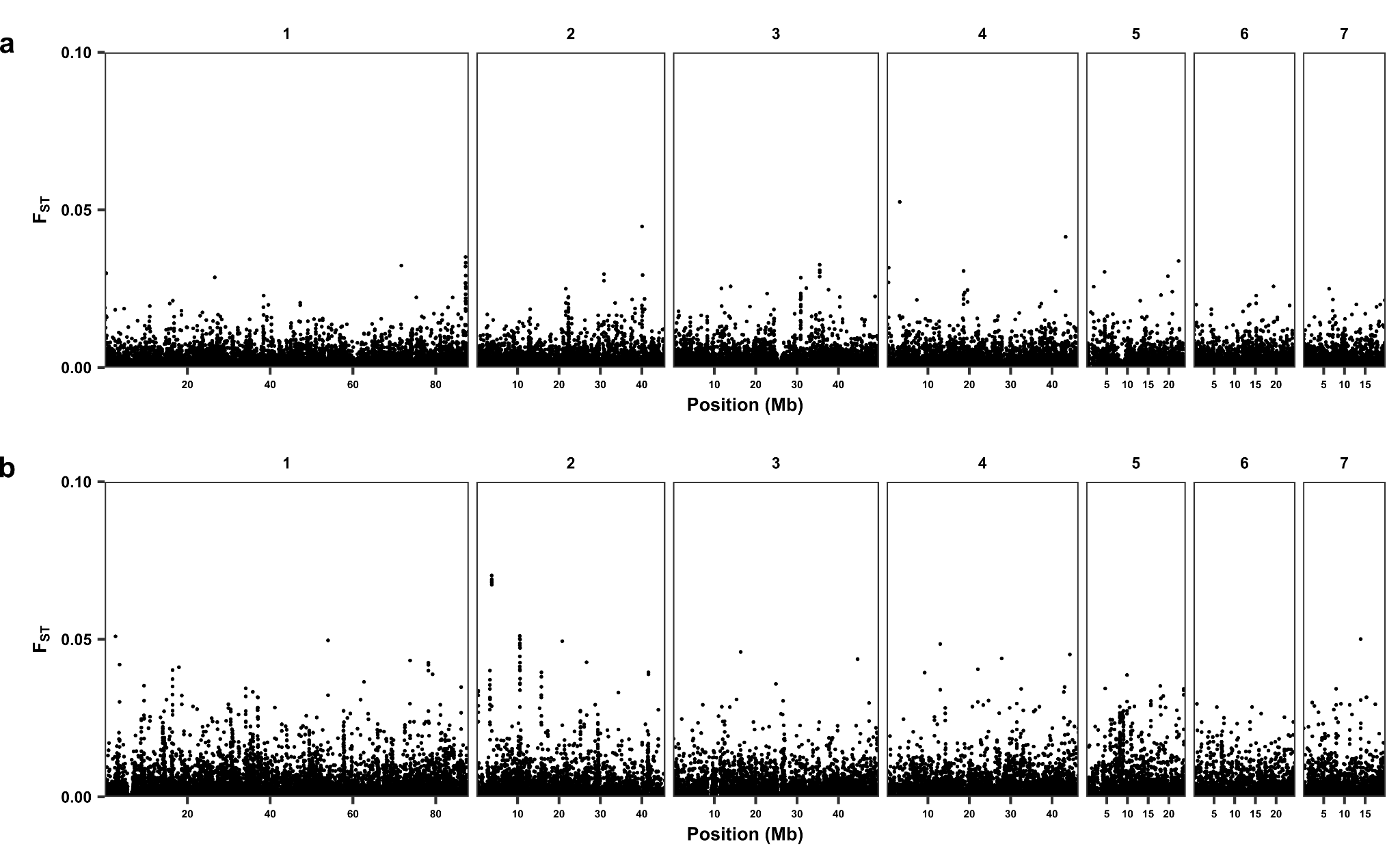

**Supplementary Figure 11. Genome-wide fixation index (*F*_ST_) values between pre- and post-treatment populations**. Values were calculated between **a)** Southern Ugandan populations (*n* = 201 pre-treatment, *n* = 127 post-treatment) or **b)** Koome group island populations (*n* = 123 pre-treatment, *n* = 51 post-treatment) in 5 kb non-overlapping windows along each autosome. Points represent median *F*_ST_ values for each window.

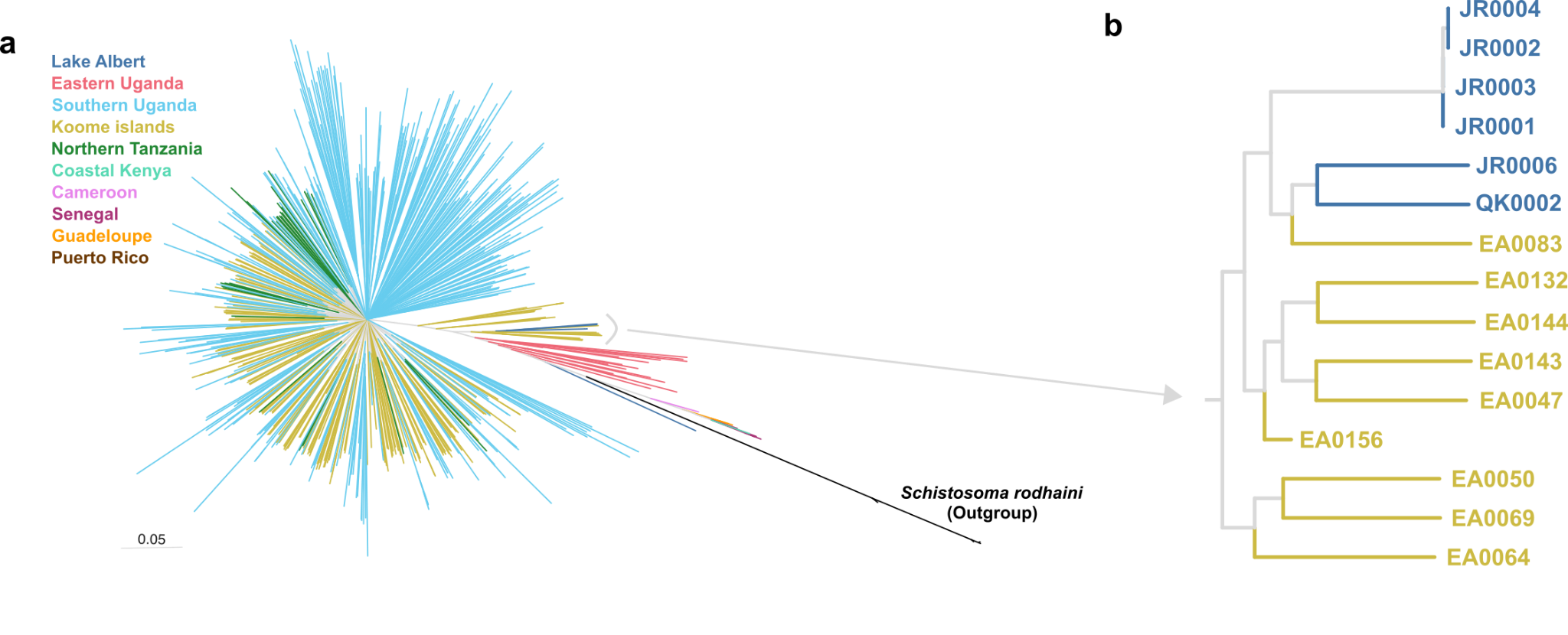

**Supplementary Figure 12. a)** Maximum-likelihood phylogenetic tree inferred using 188,923 autosomal single-nucleotide polymorphisms (SNPs) and all 574 accessions. Phylogenetic inference was conducted using IQ-TREE using a best-fit substitution model with ascertainment bias correction. Branches are coloured based on the geographical region they were sampled from and the tree is rooted on *S. rodhaini*. **b)** Highlighted clade containing samples potentially imported to the Koome group islands from Lake Albert.

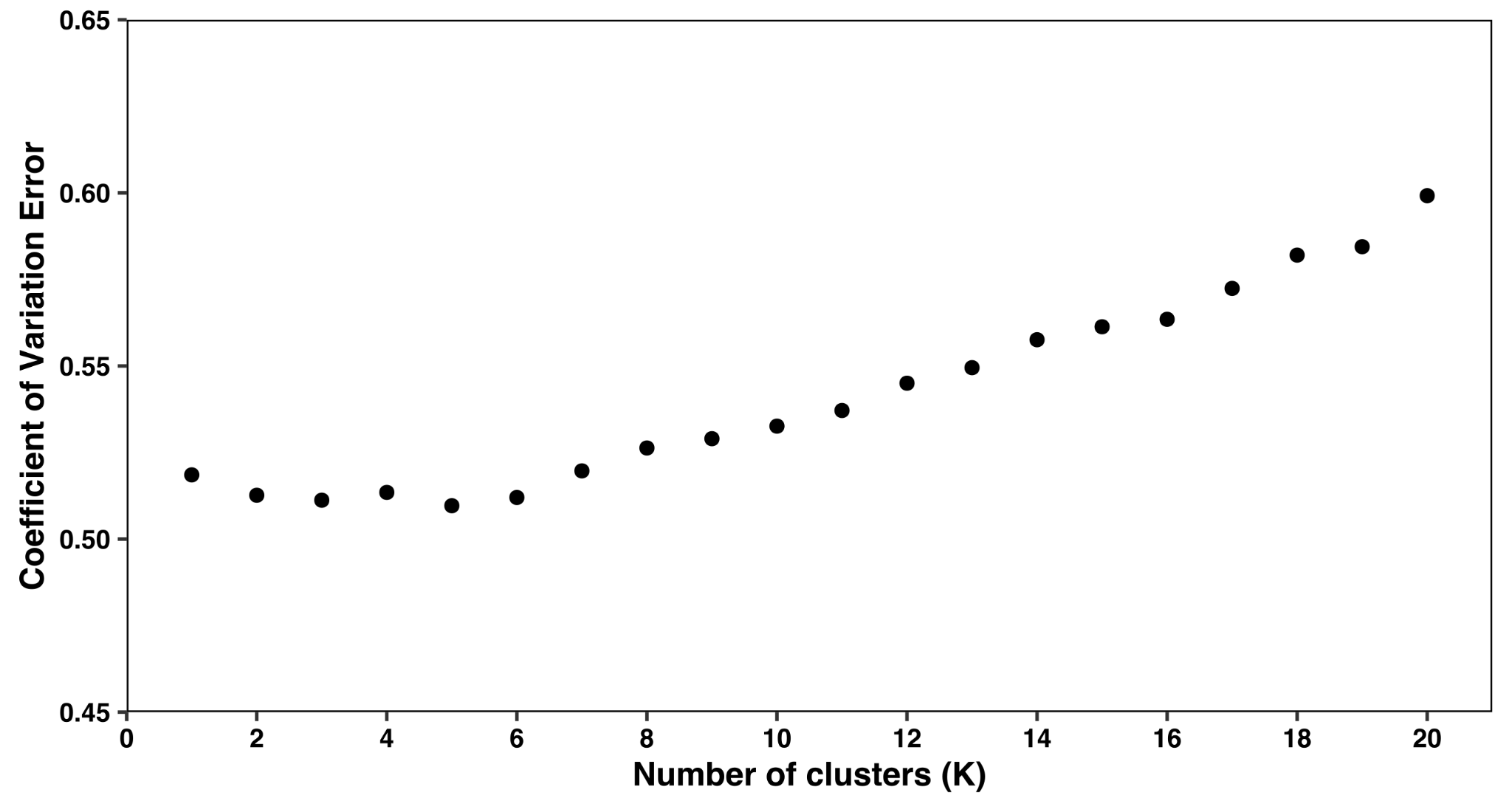

**Supplementary Figure 13.** The coefficient of variation (CV) values generated by ADMIXTURE with K values ranging from 1 to 20, 10-fold cross-validation, and standard error estimation with 250 bootstraps are shown. CV scores are shown for each K value.

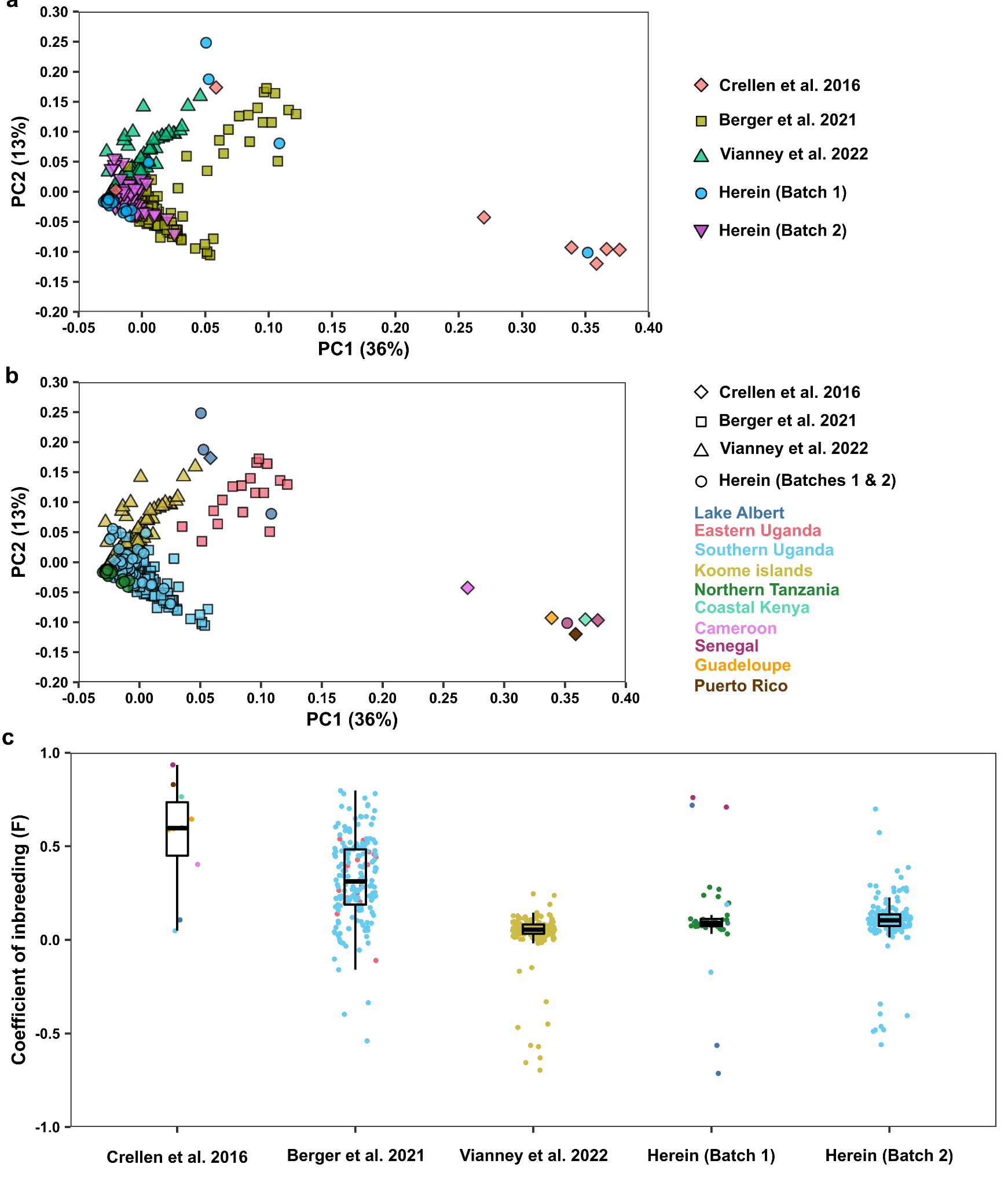

**Supplementary Figure 14.** Principal component analysis (PCA) of genetic differentiation between 505 unrelated *S. mansoni* accessions using 214,445 autosomal SNPs. Points are coloured and shaped by a) publication or b) by geographical origin and publication. c) Coefficient of inbreeding (F), points represent values for each accession, grouped by publication (x-axis) and coloured by sampling location as in b). Batch 1: Samples with identifier prefix JR*, GN* or FS*, Batch 2: Samples with identifier prefix MK*.

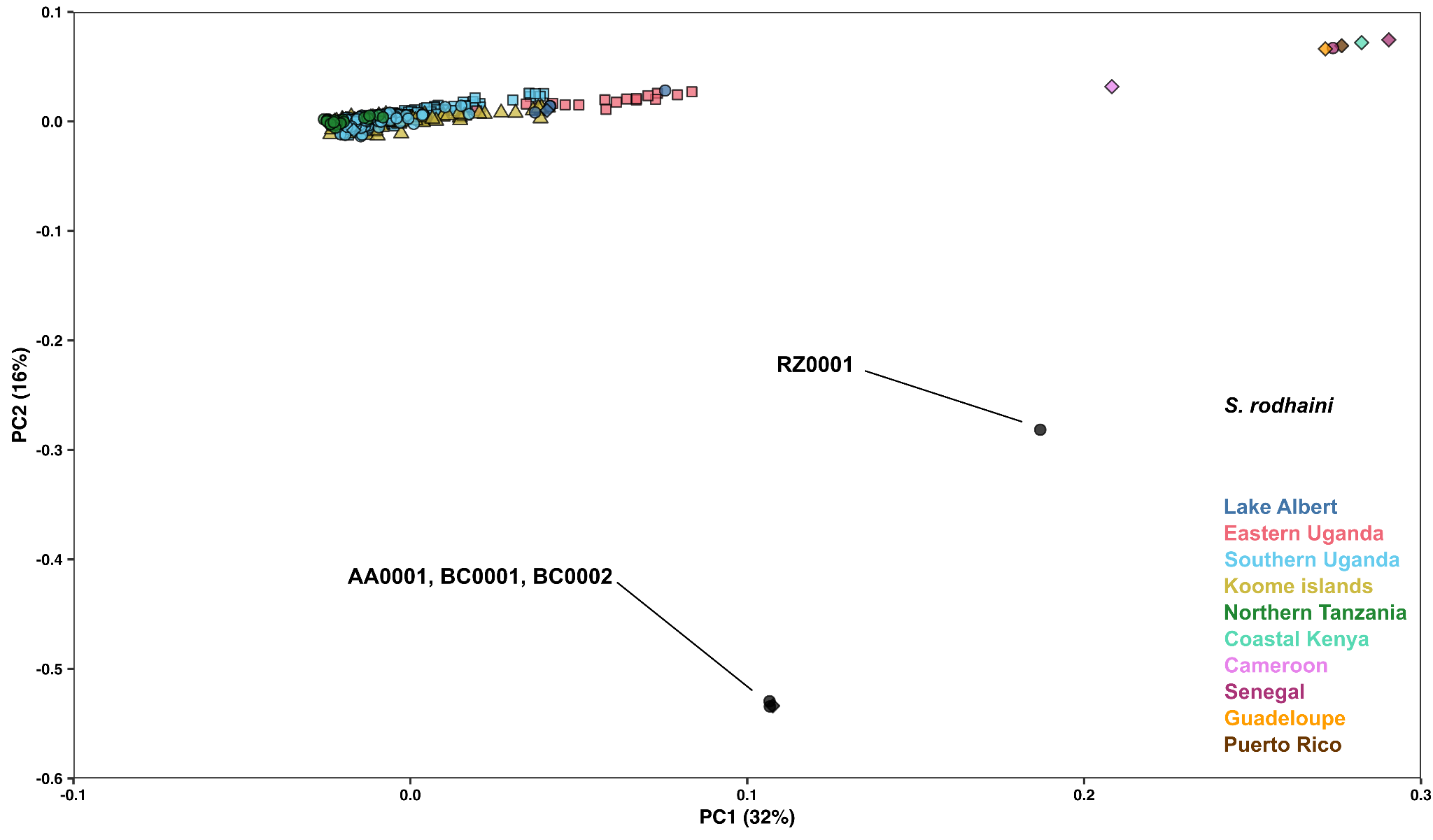

**Supplementary Figure 15.** Principal component analysis (PCA) of genetic differentiation between 505 unrelated *S. mansoni* (*n* = 542), *S. rodhaini* (*n* = 3; AA0001, BC0001, BC0002) or *S. mansoni*-*S. rodhaini* hybrid (*n* = 1; RZ0001) accessions using 201,213 autosomal SNPs. Points are coloured by geographical origin, point shapes represent the original publication from which each accession is derived: Crellen et al. (diamond), Berger et al. (square), Vianney et al. (triangle), published herein (circle).
